# TRPV1 Opening is Stabilized Equally by Its Four Subunits

**DOI:** 10.1101/2023.01.26.525787

**Authors:** Shisheng Li, Phuong Tran Nguyen, Simon Vu, Vladimir Yarov-Yarovoy, Jie Zheng

## Abstract

Capsaicin receptor TRPV1 is a nociceptor for vanilloid molecules such as capsaicin and resiniferatoxin (RTX). Even though cryo-EM structures of TRPV1 in complex with these molecules are available, how their binding energetically favors the open conformation is not known. Here we report an approach to control the number of bound RTX molecules (0-to-4) in functional mouse TRPV1. The approach allowed direct measurements of each of the intermediate open states under equilibrium conditions at both macroscopic and single-molecule levels. We found that RTX binding to each of the four subunits contributes virtually the same activation energy, which we estimated to be 1.86 kcal/mol and found to arise predominately from destabilizing the closed conformation. We further showed that sequential bindings of RTX increase open probability without altering single-channel conductance, confirming that there is likely a single open-pore conformation for TRPV1 activated by RTX.

## Introduction

Allosteric coupling links structurally separate function domains together through a global conformational rearrangement which, in protein complexes, introduces cooperativity (1). Cooperative activation can substantially enhance the sensitivity to a stimulus, hence bestows a significant functional advantage to multi-subunit complexes. Indeed, multi-subunit complex formation is a common phenomenon in biology (2). Even though diverse models have been satisfactorily used to describe the overall cooperative behavior in protein complexes (3, 4), it remains a major challenge to directly access cooperative activation among subunits as it occurs, limiting our understanding of the underlying mechanism. The Monod-Wyman-Changeux (MWC) model proposed over half a century ago (3) postulates that ligand inding to individual subunits promotes a concerted activation transition by an identical cooperative factor, *f*, which reflects energetic contribution by each subunit (Fig. 1A). However, exactly because of cooperative activation, the intermediate states (shaded in Fig. 1A) are rapidly traversed; their energetic contributions are rarely determined individually. Reflecting this challenge, a common practice has been to fit observed activities to a Hill function, yielding a slope factor estimate that is neither the cooperative factor nor the number of subunits but their combined contribution (5).

**Figure 1.**
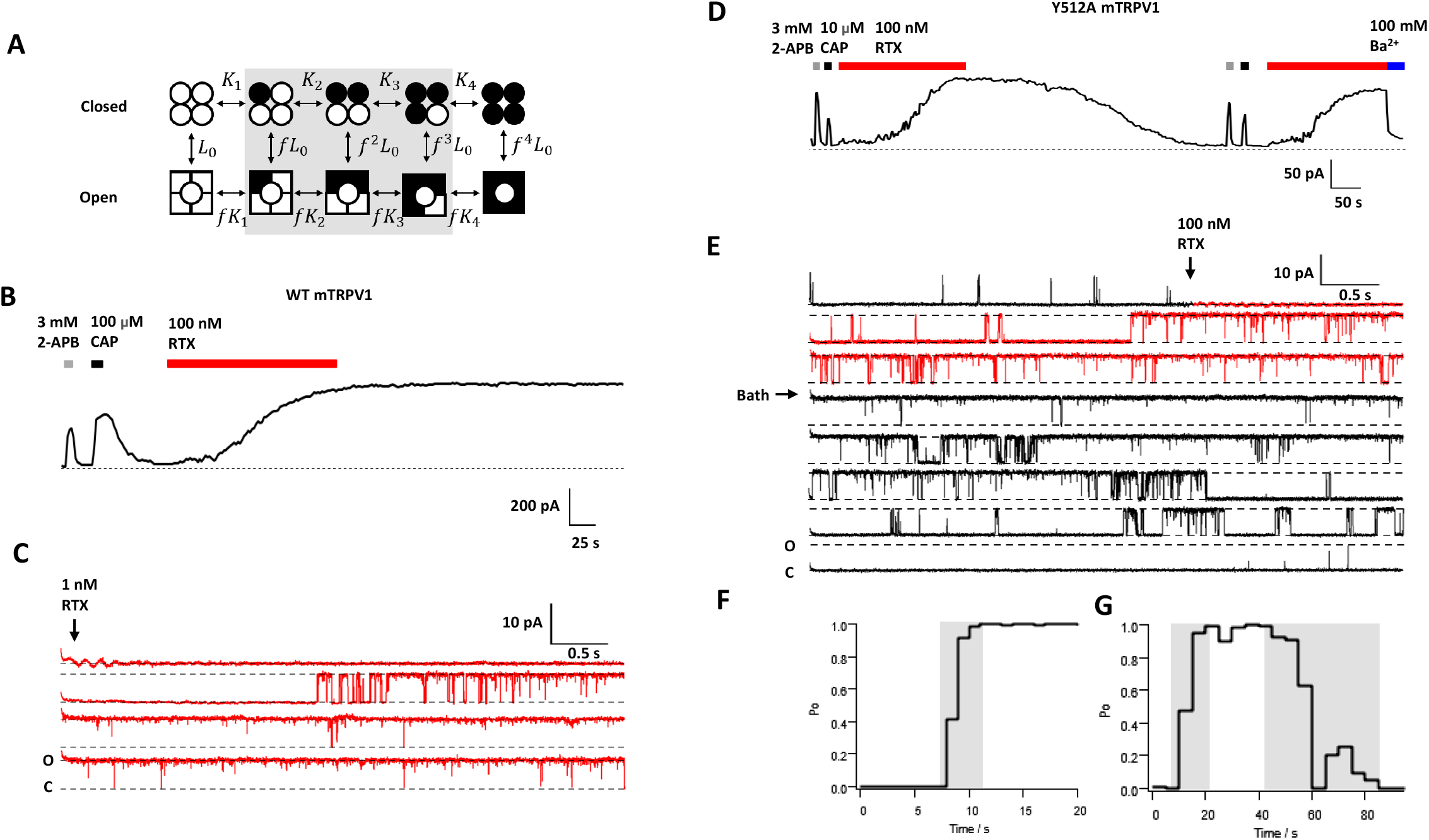
Intermediate open states of wildtype and Y512A mutant channels. A: A classic MWC-type allosteric model for TRPV1 activation; circles and squares indicate a resting and activated subunit, respectively; open and filled symbols represent apo and RTX-bound subunits, respectively. *L*_*0*_, *K*_*n*_, *f*^*n*^*K*_*n*_ and *f*^*n*^*L*_*0*_ are equilibrium constants for each transition (n = 1 to 4). B: A representative macroscopic inside-out patch-clamp current trace of wildtype mTRPV1 at +80 mV. C: A representative single-channel recording of wildtype mTRPV1 activated by RTX. D: A representative macroscopic current trace of Y512A. Ba^2+^ was used to block channel current. E: A representative single-channel recording of Y512A activated by RTX. F&G: Po versus time plots for the recording shown in C and E, respectively. Transitional Po phases are highlighted by grey shading. Red segments in C and E indicate duration of RTX application.

Biological ion channels are usually made of multiple, functionally coupled subunits or repeating domains (6, 7). The capsaicin (CAP) receptor TRPV1 is a representative multi-subunit ion channel (8, 9) whose activation exhibits allosteric properties (10-13). In a typical recording of the mouse TRPV1 activated by CAP, current amplitude reached over 90% maximum within hundreds of milliseconds (Fig. S1). Even though this activation process is slow among ion channels, the intermediate states were swiftly traversed and could not be individually identified and studied (14). Methods to slow down activation and to fix gating transition in individual subunits were developed in this study, which allowed direct assessment of the property of intermediate states and subunit coupling at equilibrium under physiological conditions.

## Results

We first used resiniferatoxin (RTX), a potent TRPV1 activator (8), to replace CAP as the agonist. RTX binds in the same vanilloid binding pocket as CAP (9) but activates the channel with an extremely slow (in minutes) time course; RTX-induced activation is also irreversible (Fig. 1B). The slow activation time course revealed a clear transition phase of increasing open probability in single-channel recordings (Fig. 1C and 1F), indicating sojourns in unstable intermediate open states as suggested by the MWC model. However, these transient and stochastic events were observed under non-equilibrium conditions, hence were still difficult to analyze.

From the RTX-TRPV1 complex structures (9, 15), we identified key channel residues that directly interact with a bound RTX (Fig. S2). Mutations to these residues in most cases either exerted minor gating effects or eliminated function (Fig. S3). Mutating Y512 to alanine retained the slow activation property of the wild-type channel but noticeably made RTX activation completely reversible (Fig. 1D). We have previously suggested that Y512 serves to physically block a bound vanilloid molecule from exerting the binding pocket (16), in agreement with structural observations (15). Molecular docking of RTX to TRPV1 structures in the closed (C1 and C2) and open (O1) states using RosettaLigand (17, 18) suggested reductions in Rosetta binding energy by this point mutation, in supportive of weakening of RTX binding (Fig. S4). After washing off RTX, Y512A channels could be activated by 2-APB, CAP and RTX similar to naïve channels without detectable desensitization (under the Ca^2+^-free condition). Single-channel recordings confirmed that Y512A traversed the transition phase during both activation and deactivation time courses (Fig. 1E&G; see also Fig. S5).

Taking advantage of the reversibility property of the Y512A mutant, we studied rat TRPV1 concatemers containing different numbers of wildtype and Y511A mutant (equivalent to Y512A in mouse TRPV1) subunits (Fig. 2A). These concatemer channels were previously shown to respond to CAP in a concentration-dependent manner, with the CAP potency progressively decreased as the number of mutant protomers increased (19). We confirmed that the spontaneous open probability of each concatemer matched that of wildtype channels (Fig. 2D&E). We observed with macroscopic (Fig. 2B) and single-channel recordings (Fig. S6) that RTX could fully activate all concatemer channels. Importantly, currents from these concatemers exhibited gradually enhanced reversibility upon washing off RTX, with channels containing all or three wildtype subunits (YYYY and YYYA, respectively) exhibiting little current deactivation, whereas those with all Y511A subunits (AAAA) exhibiting complete deactivation like channels formed by unlinked monomeric Y511A subunits (see Fig. 1D). We further confirmed that the remaining currents were not due to an unstable patch (Fig. 2B) but instead represented steady intermediate open probabilities (Fig. 2C; see also Fig. S6). These behaviors are expected as RTX binding to the Y511A subunits in concatemers is reversible, but irreversible when bound to the wildtype subunits. Therefore, after extended washing, the concatemer channels contained a fixed number (zero to four) of bound RTX molecules.

**Figure 2.**
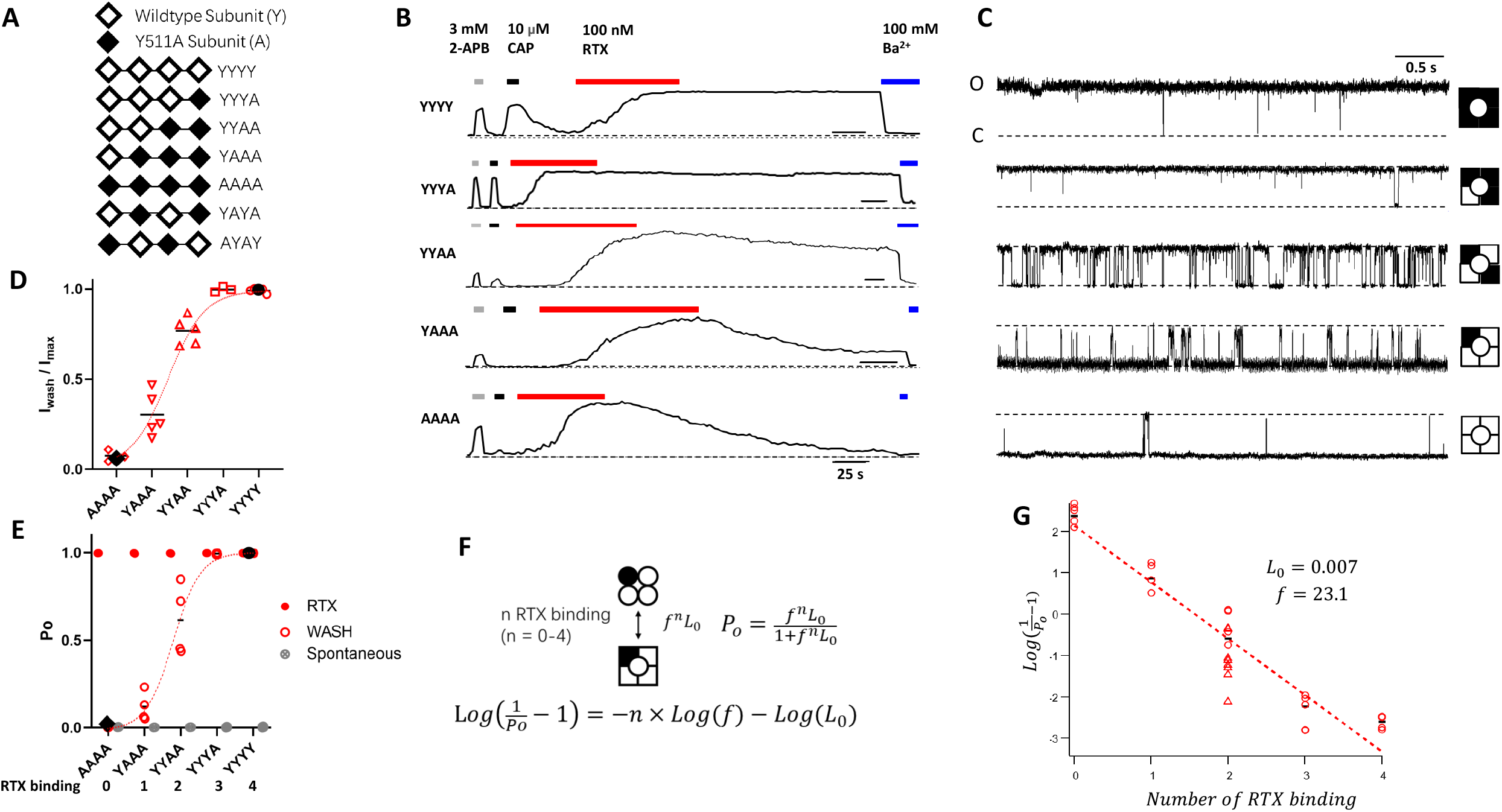
Equilibrium measurements of transition states. A: Illustration of concatemer cDNA constructs. B&C: Representative current traces from concatemer channels. RTX activation reached near 100% Po and was partially or fully reversible, depending on the number of mutant subunits. The Y511A mutation did not affect 2-APB activation and, as expected, shifted the concentration dependence for capsaicin activation to the right (16, 19). D: Normalized remaining currents after complete wash (Iwash/Imax) can be well-fitted to the function *Po=(f*^*n*^*\*L*_*0*_*)/(1+f*^*n*^*\*L*_*0*_). E: Spontaneous (filled grey circles), maximal (filled red circles) and remaining Po (open red circles) measurements from single-channel recordings fitted to the same function as in D. For D&E, black diamonds and circles represent the level of the spontaneous activity of the apo state and maximal activity of the fully RTX-bound state, respectively, of the wildtype channel. F: The simplified MWC model when the number of bound RTX is fixed. G: Estimation of *Lo* and *f* using the model in F. Data points from YAYA and AYAY are presented in red triangles.

The concatemers allowed us to characterize each vertical equilibrium of the MWC model (Fig. 1A) in isolation. For example, after fully activating the YYAA channels and washing off RTX from the mutant subunits, each channel contained two RTX molecules (the third column of the model). The open probability (Po) value of these channels could be determined from both macroscopic currents (77 ± 3% of the maximum response; Fig. 2D) and single-channel currents (62 ± 10%; Fig. 2E) at thesteady state. Po estimates for YYYY and AAAA channels matched those fromchannels made of wildtype and mutant monomeric subunits, respectively (Fig. 2D&E). The Po measurements from all concatemers could be satisfactorily fitted to a general formula derived from the MWC model that describes the relationship between Po and the number of bound ligands (Fig. 2F). Transformation of the formula yields a linear function that allows easy estimation of the equilibrium constant of unliganded channels, *L*_*0*_, and the cooperative factor, *f*. Fig. 2G shows results from the single-channel data. We estimated the *L*_*0*_ value to be 0.007, equivalent to a resting Po of 0.7%, which is consistent with experimental observations from wildtype channels (20); we estimated the *f* value to be 23.1, equivalent to an activation energy of 1.86 kcal/mol per RTX binding (Fig. 2G).

It is noticed that a linear fit agreed with experimental measurements remarkably well, suggesting that subunit contributions to activation are indeed equal, as postulated by the MWC theorem. The position of wildtype and mutant subunits within the concatemers had no obvious influence on the activation energy, as data from concatemers YAYA and AYAY agreed reasonably well with that from YYAA (Fig. 2G), an observation consistent with the new cryo-EM data showing that RTX binding has no position preference (15).

Where does the 1.86 kcal/mol activation energy arise from? For the isolated transition shown in Fig. 2F, the equilibrium can be altered at either the closed or the open state, both of which could contribute to an increased equilibrium constant. To quantify energetic effects of RTX binding to the closed and open state, we analyzed single-channel dwell-time distributions of the concatemers at steady state (Fig. 3A-C). When results from all concatemers were compared, we found that the mean dwell-time of, the closed state decreased exponentially with an increasing number of bound RTX, whereas the mean dwell-time of the open state increased exponentially (Fig. 3D). Equilibrium constant values estimated from dwell-time measurements matched closely with those estimated from open probability measurements (Fig. 3E). From linear fittings of the log-transformed dwell-times (Fig. 3D), we estimated that each RTX binding destabilized the closed state by 1.17 kcal/mol, stabilized the open state by 0.53 kcal/mol. Therefore, activation of TRPV1 by RTX originates predominately from destabilization of the closed state.

**Figure 3.**
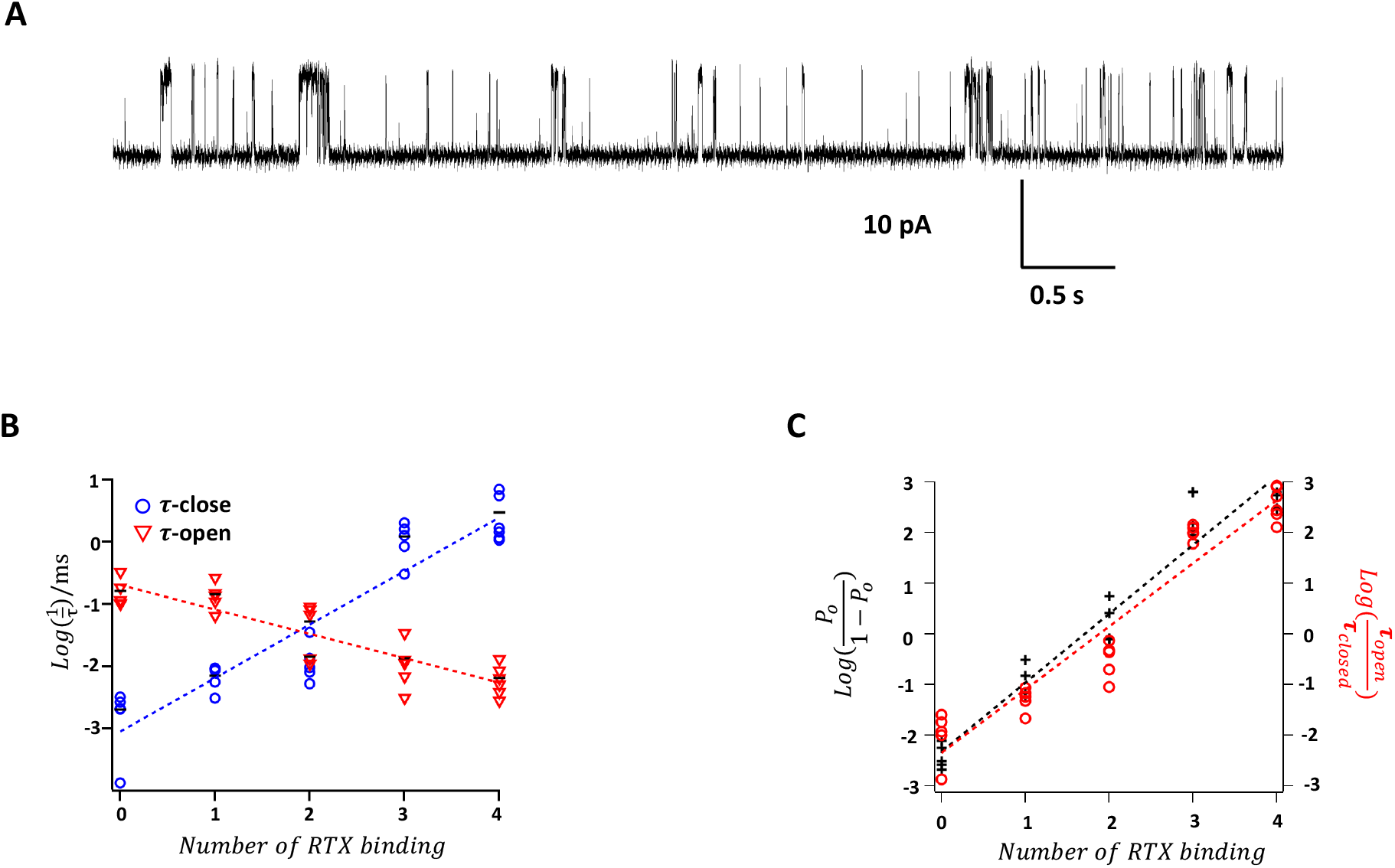
Single-channel dwell-time analysis reveals energetic effects on the closed and open states. A. An example single-channel trace from the YAAA channel. B. Relationship between dwell-time in the closed or open state and the number of RTX binding. Dotted lines represent linear fits of the log-transformed data, with a slope factor of 0.86 for the closed state and -0.39 for the open state. The mean values are shown as black bars. n = 3-5. C. Comparison between log-transformed lifetime ratios (red) and open probabilities (black). Dash lines represent linear fits, with slope factor values of 1.36 (Po) and 1.25 (lifetime ratio).

Measurements from the concatemers revealed the Po values with 0-to-4 bound RTX molecules. Using this information, we could tentatively identify the intermediate open states produced by sequential bindings of RTX during the course of activation. A typical time course of the wildtype channel single-channel activation exhibited distinct open states with low, moderate, and high Po values, corresponding to one, two, and three or four bound RTX molecules, respectively (Fig. 4A). All these open states, observed in the same single-channel recording, exhibited identical amplitudes M(Fig. 4B&C). Similar observations were made from the concatemers. This finding is consistent with the classic MWC model in which the multi-subunit complex exists in two (tense and relaxed) conformations, with ligand binding progressively shifting the equilibrium towards the relaxed, a.k.a. open, conformation (3). Our results suggest that TRPV1 activated by RTX has a single open pore conformation.

**Figure 4.**
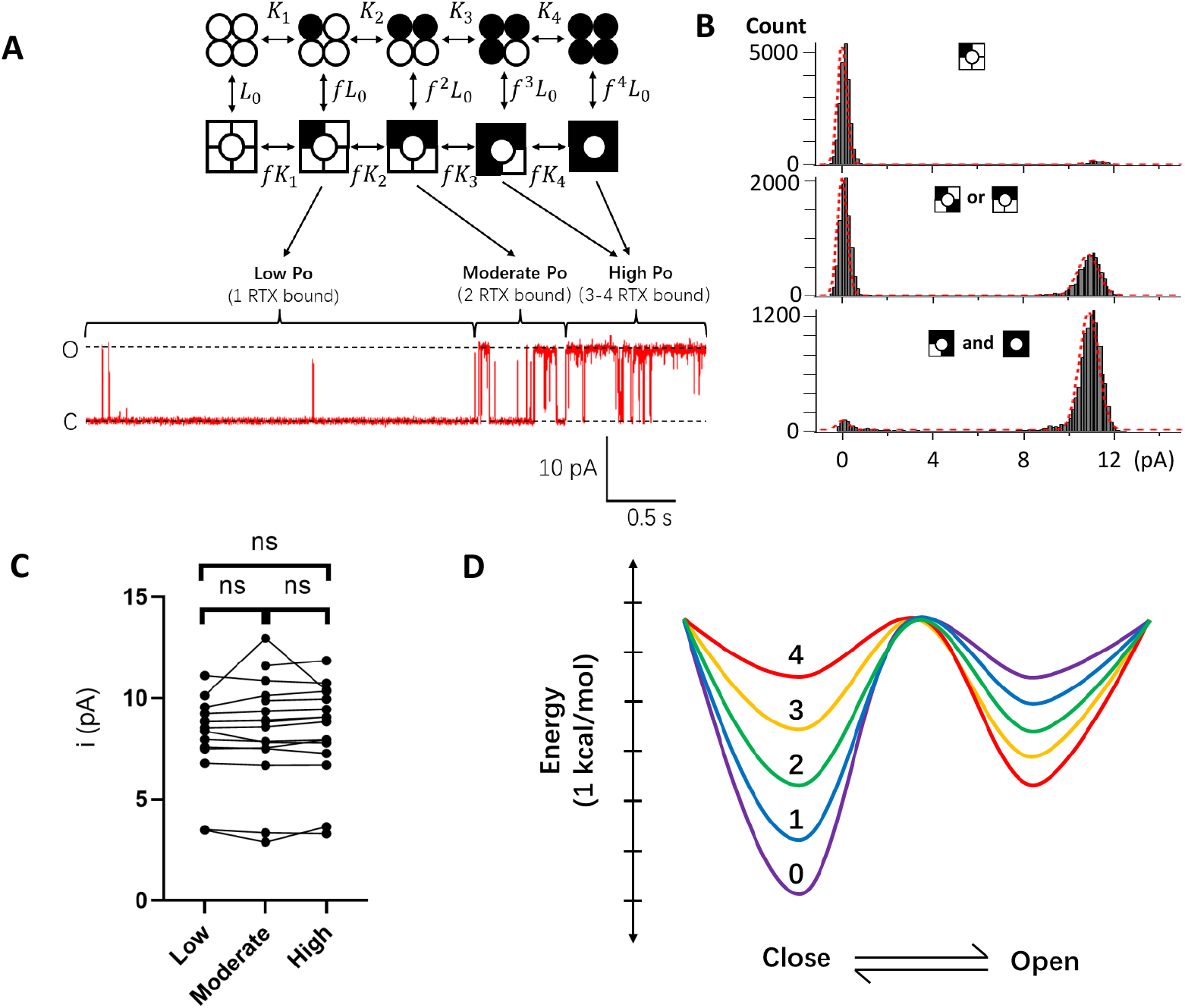
RTX-induced open states exhibit identical single-channel conductance. A: A representative single-channel trace from the wildtype TRPV1 activated by RTX. Distinct states with varying numbers of bound RTX were assigned according to Po. B: All-point histograms (with double-Gaussian fitting) of the three phases shown in panel A; only the open probabilities, represented by the relative areas of the two Gaussian functions, changed but not the single-channel amplitude. C. Comparison of single-channel amplitudes of distinct phases measured from 16 individual patches; ns, no significant difference. D. Eyring energy plots for channels with varying numbers of bound RTX. Ligand binding shifts the closed state energy more substantially than the open state; the four ligand binding steps shift these energies by the same amount; the energy wells for the closed and open states remain at the same locations along the reaction coordinate. The level of energy barrier is artificially set and unchanged.

## Discussion

Previously we reported that structurally related natural vanilloids from chili peppers and gingers as well as synthetic capsaicin derivatives induce similar conformational changes of the vanilloid binding pocket of TRPV1, even though they may bind in different poses (21-23). We now present data suggesting that the, activated channel pore adopts the same open conformation when a varying number of vanilloid binding pockets are occupied by RTX, consistent of pore opening being a concerted transition. Putting together, these observations support the general assumption that TRPV1 behaves as an allosteric protein (10-13). By fixing the number of bound RTX molecules, we were able to directly quantify equilibrium properties from otherwise unstable intermediate states at both macroscopic and single-molecule resolutions. We observed binding of RTX to each of the four subunits yielded an equal activation energy, as predicted by the classic MWC model for homotropic cooperativity. A recent structural study of RTX-TRPV1 complexes indeed suggested non-preference in RTX binding to the four subunits (15). The activation energy, estimated at 1.86 kcal/mol per liganded subunit, is comparable to that of other channels such as CNG channels (0.84-to-1.3 kcal/mol) determined indirectly from overall ligand activation behaviors (24, 25). The comparable maximal Po of capsaicin-activated channels indicates that capsaicin binding is expected to produce a similar activation energy (16). Combining the activation energies from multiple subunits contributes to an exponential increase in open probability (the 1.86 × 4 = 7.44 kcal/mol activation energy shifts the closed-to-open equilibrium toward the open state by over 280000 folds) (Fig. 4D). For TRPV1 activated by RTX, the shift in equilibrium originates mostly from destabilization of the closed state.

Energetic contributions from the four subunits represent substantial activation cooperativity in TRPV1. Indeed, cooperative activation is a common phenomenon for diverse allosteric proteins. However, a general challenge for studying a cooperative process is to capture the intermediate states with either functional or structural methods. In the absence of direct information concerning the intermediate states, a mechanistic interpretation is often speculative. The Monod-Wyman-Changeux (MWC) model for allosteric proteins made of identical subunits (3) has been successfully applied to a wide variety of proteins including those made of different subunits. Nonetheless, equivalence in subunit contributions, a key feature distinguishing it from a Koshland-Némethy-Filmer (KNF)-type sequential model(26), remains a postulation. One key validation of the MWC theorem—measuring directly from the intermediate activation states—has been lacking, due to cooperativity among subunits that makes transitional states intrinsically unstable. Here we designed methods locking TRPV1 in each of the intermediate activation states with varying ligand occupations, which allowed direct measurements of their thermodynamic properties under physiological conditions at equilibrium. Near-perfect equivalence in subunit contribution to allosteric coupling was confirmed.

All RTX-induced open states are found to produce identical single-channel current amplitudes, suggesting the activated pore adopts near identical conformations. Therefore, pore opening by RTX appears to be driven by a concerted conformational change. Our functional data hence suggest that structural asymmetry of the activated states occurs mainly in the vanilloid binding pockets, consistent with the recently eported RTX-TRPV1 complex structures (15). Based on the 1.86 kcal/mol/subunit activation energy, channels with 1-to-4 bound RTX molecules are expected to exhibit an open probability of 13.9%, 78.9%, 98.8% and 99.9%, respectively. Channels with two bound RTX molecules already spend two-third of the time in the open state under physiological conditions. Only the fully RTX-bound channels exhibited an open pore conformation in cryo-EM studies (15, 27), suggesting a shift in stability of the closed and/or open conformation under these experimental conditions, such as lower temperatures (20).

Concerted pore opening has been previously proposed for other channel types including Kv channels, for which independent voltage-sensor movements in the four subunits precede channel activation (28, 29). Very brief (microseconds) subconductance states were found to be associated with asymmetrical subunit activations by voltage, hence representing intermediate gating states (30, 31). It was proposed that subconductance states could arise from filtering effects on very rapid flickering events—representing likely a concerted transition—between the fully open state and the closed state, with voltage-dependent subunit activations progressively shifting the equilibrium towards the open state (30). While TRPV1 activation by RTX exhibited similar progressive shifting towards the open state, dwell-times in the closed and open states appeared to be long enough to skip the filtering effect on current amplitude. The allosteric behavior described in the present study may help correlate published high-resolution channel structures (15, 32) with distinct functional states towards a better understanding of the activation mechanism of TRPV1 and other ion channels.

## Materials and Methods

### Cell culture and molecular biology

The mouse TRPV1 wildtype and Y512A mutant cDNAs were constructed into pEYFP-N3 vector, where the C-terminus of the channel was fused with the Cdna encoding enhanced yellow fluorescent protein (eYFP) to help to identify transfected cells during patch clamp experiments, as previously described (33). The Y512A point mutation was introduced using a mutagenesis kit (Agilent Technologies) and confirmed by sequencing. Concatemers YYYY, YYYA, YYAA, YAYA, AYAY, YAAA, and AAAA represent tandem tetrameric cDNA constructs of rat TRPV1 wildtype (Y) and Y511A (A, equivalent to Y512A in mouse channel), were generous gifts from Dr. Avi Priel and have been previously described (19).

HEK293T cells (purchased from American Type Culture Collection) were cultured in a DMEM medium (Hyclone or Gibco) supplemented with 10% (v/v) fetal bovine serum (FBS) (Corning or GenClone), 1% (v/v) penicillin/streptomycin (Fisher Scientific) and 1% (v/v) MEM Non-Essential Amino Acids Solution (Hyclone) at 37°C with 5% CO_2_. Cells were cultured onto 25 mm glass coverslips (Fisher Scientific) in 35 mm chambers 18-24 h before transfection. Transient transfection was conducted 18-24 h before patch-clamp recording using Lipofectamine 2000 (Invitrogen) according to the manufacturer’s instructions. For macroscopic recordings, 1 μg plasmid was used for transfection for each chamber; for single-channel recordings, 0.1 μg plasmid was used. For concatemers channels without a fluorescent tag, the YFP plasmid was co-transfected (0.2 μg) for each chamber.

### Electrophysiology

Patch-clamp recordings were done using an EPC10 amplifier controlled with PatchMaster software (HEKA) in configurations as specified in the Results. Patch pipettes were pulled from borosilicate glass (A-M systems or Sutter Instrument) using Sutter Instrument P-97 micropipette puller and fire-polished to 2-to-6 MΩ for macroscopic recordings, or 8-to-15 MΩ for single-channel recordings. For whole-cell recordings, serial resistance was compensated by 60%. Current signal was filtered at 2.3-to-2.9 kHz and sampled at 10.0-to-12.5 kHz. All recordings were conducted at room temperature. Ruthenium red (10 μM) and/or Ba^2+^ (100 mM) were used to block channel current for checking leak. A holding potential of 0 mV was used, from which a 300-ms step to +80 mV followed by a 300-ms step to -80 mV was applied. The durations of the +80 mV and -80 mV steps were between 300 ms and 2 s, adjusted according to experimental needs. The voltage was held at +80 mV for long continuous single-channel recordings. Standard symmetrical bath and pipette solutions contained (mM) 140 NaCl, 0.2 EGTA, 10 glucose (optional) and 15 HEPES (pH 7.2-to-7.4). Solution switching was achieved with a Rapid Solution Changer (RSC-200, Biological Science Instruments). 2-APB was dissolved in DMSO to make 1 M stock solution and diluted to working concentrations using the bath solution; capsaicin was dissolved in DMSO to make 50 mM stock solution and diluted to working concentrations using the bath solution; RTX was dissolved in ethanol to make 1 mM stock solution and diluted to working concentrations using the bath solution.

### Rosetta modeling of RTX binding to mutant TRPV1 channels

To evaluate RTX binding to TRPV1, we used the cryoEM structures of TRPV1 solved in complex with RTX in the open (O1 state, PDB:7l2l) and closed (C1 state, PDB: 7l2n; C2 state, PDB:7mz5) states to model RTX interaction. RosettaLigand(17) was used to dock RTX to wild-type and Y511A mutant structures for each state. We generated 2,000 RTX conformers for the docking process using BioChemical Library (BCL) (18). The detail of the docking algorithm has been described elsewhere (17) (see Appendix for Rosetta docking scripts and command lines). A total of 10,000 docking models were generated and the top 1,000 models ranked by total_score were selected for analysis. Binding energy is represented by the interface_delta_X term reported in Rosetta Energy Unit (R.E.U). All-atom root-mean-square deviation (RMSD) was used to compare the generated docking RTX models to the RTX binding conformations captured in the cryoEM structures.

### Data analysis

Patch-clamp data were exported and analyzed using Igor Pro 8. Statistical analyses were done using GraphPad Prism 8. Macroscopic current amplitude was calculated by measuring the difference between ligand-activated current and the baseline current level before any ligand perfusion. A digital filter at 0.4 kHz was used for analyzing single-channel amplitude and open probability. Single-channel recordings were analyzed using all-point histograms and fitted to a double-Gaussian function; single-channel amplitude was measured as the difference between the two gaussian peaks; single-channel open probability was measured by calculating the portion of open events over the total time recorded. Only true single-channel recordings (n = 22) or two-channel recordings (n = 2) were used. Open probability of two-channel recordings was calculated using the equation 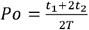, where *t*_*1*_ one-channel opening evens observed, *t*_*2*_ is the total time of two-channel opening evens observed, *T* is the total recording time. Spontaneous open probability was measured using the equation 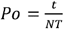, where *t* is the total time of the open events observed at room temperature in bath solution, *N* is the total number of channels in the patch which was determined by calculating the amplitude ratio between maximum current and the average single-channel current, *T* is the total recording time. There were no overlapping open events in these recordings. To analyze single channel dwell-times, the recorded single-channel traces were processed using the single-channel search function in Clampfit to generate dwell-time histograms (34).

Model fitting was conducted in Igor, using the global fitting procedure when needed. Statistical analysis was done using GraghPad Prism 8. *Student*’s t-test was used when comparing between two groups. For comparisons within the same recordings, paired t-test was used. For comparison between multiple groups, one-way ANOVA was used.

## Acknowledgements

We are grateful to Avi Priel for sharing the concatemer constructs. We thank members of the Zheng and Yarov-Yarovoy laboratories for assistance and discussion.

---

**Fig.S1.**
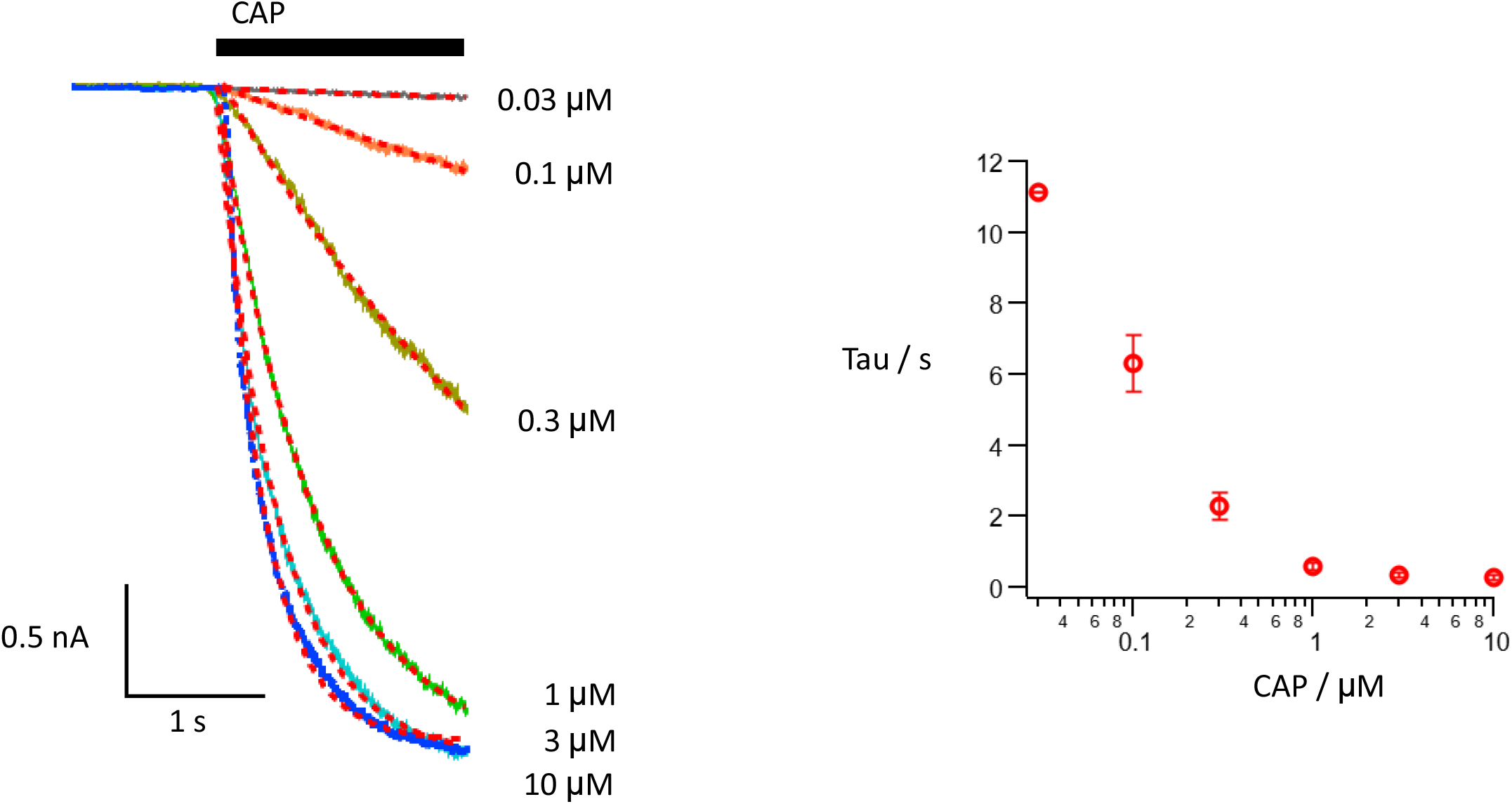

**Fig.S2.**
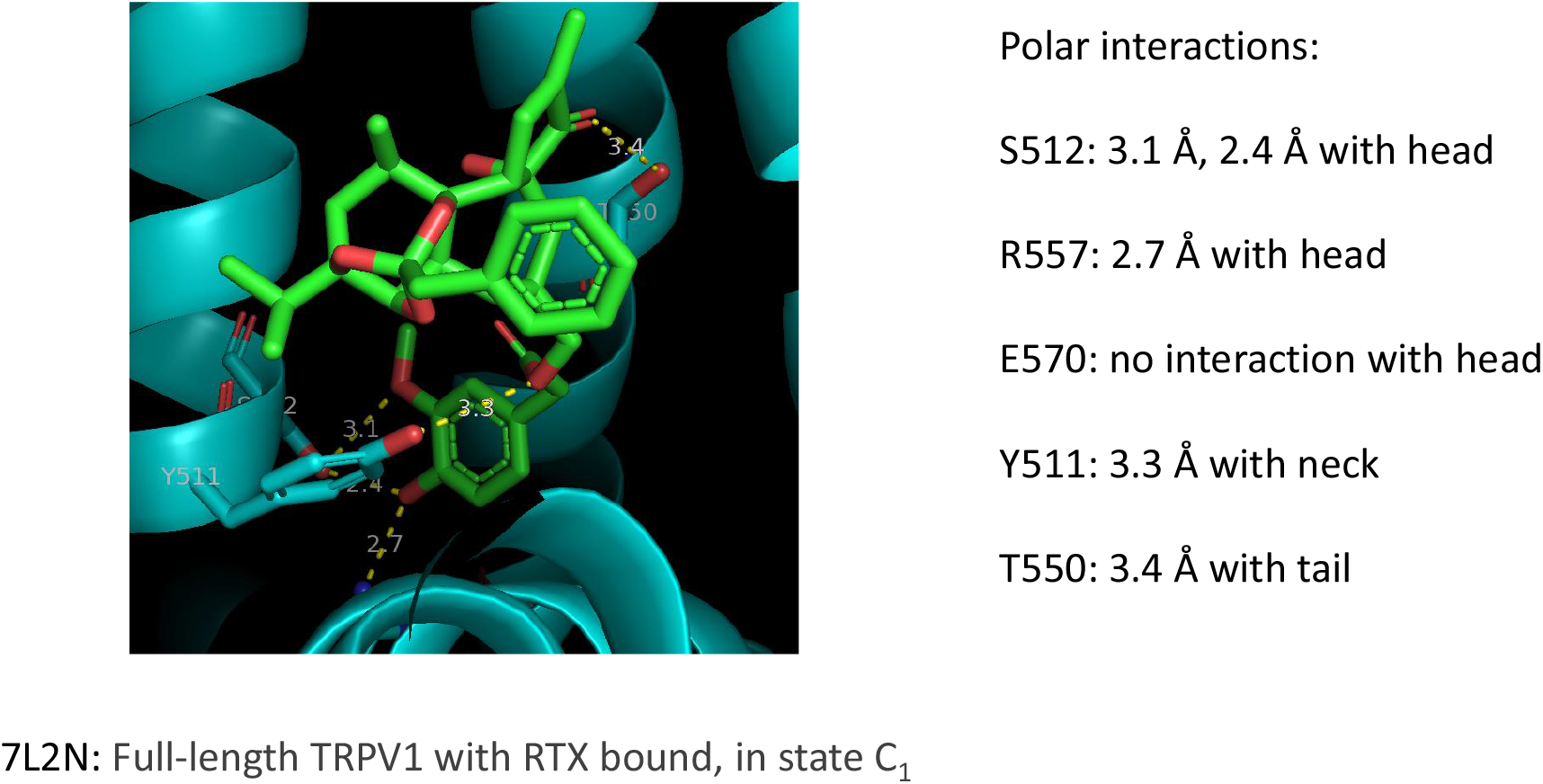
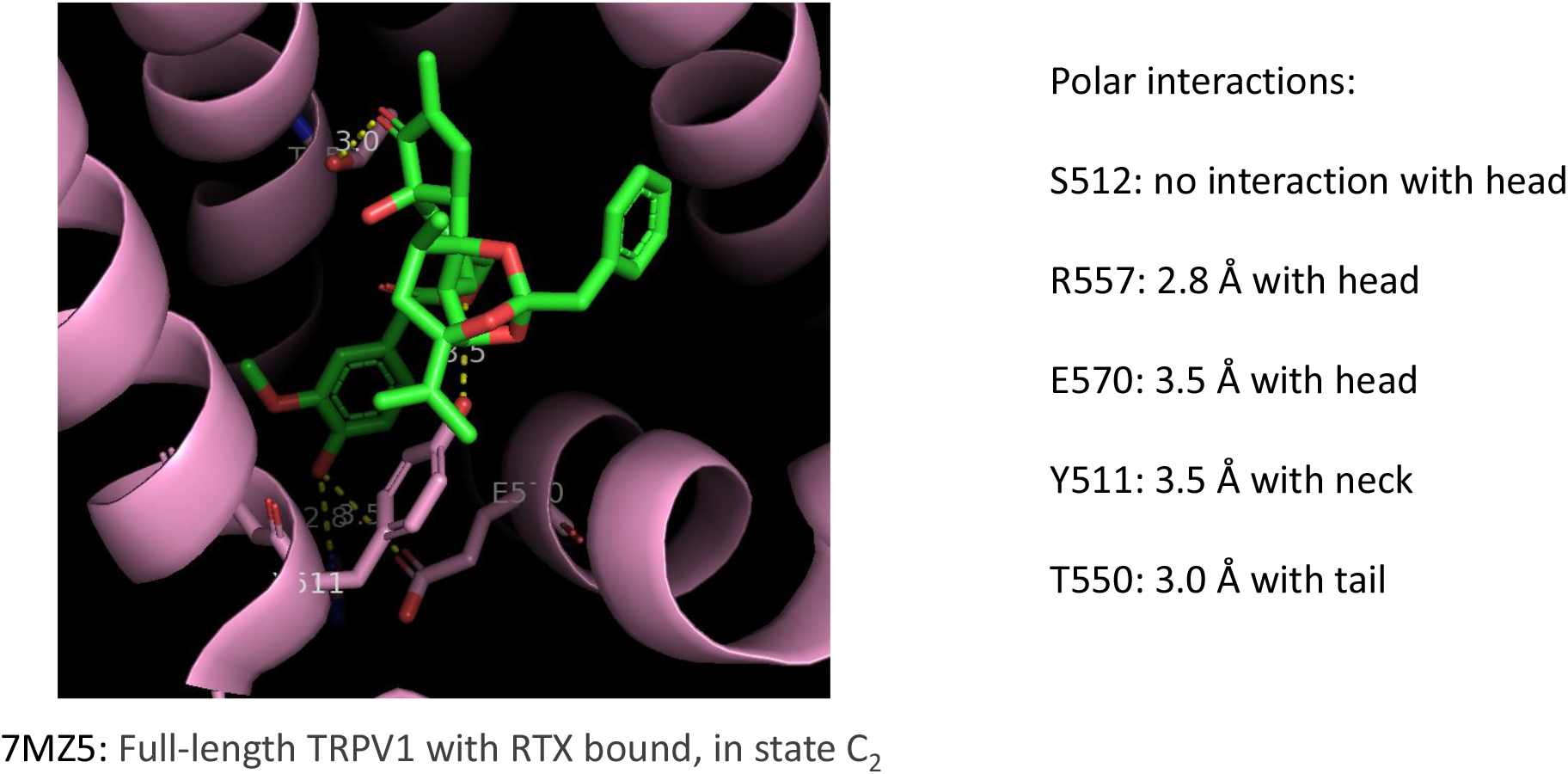
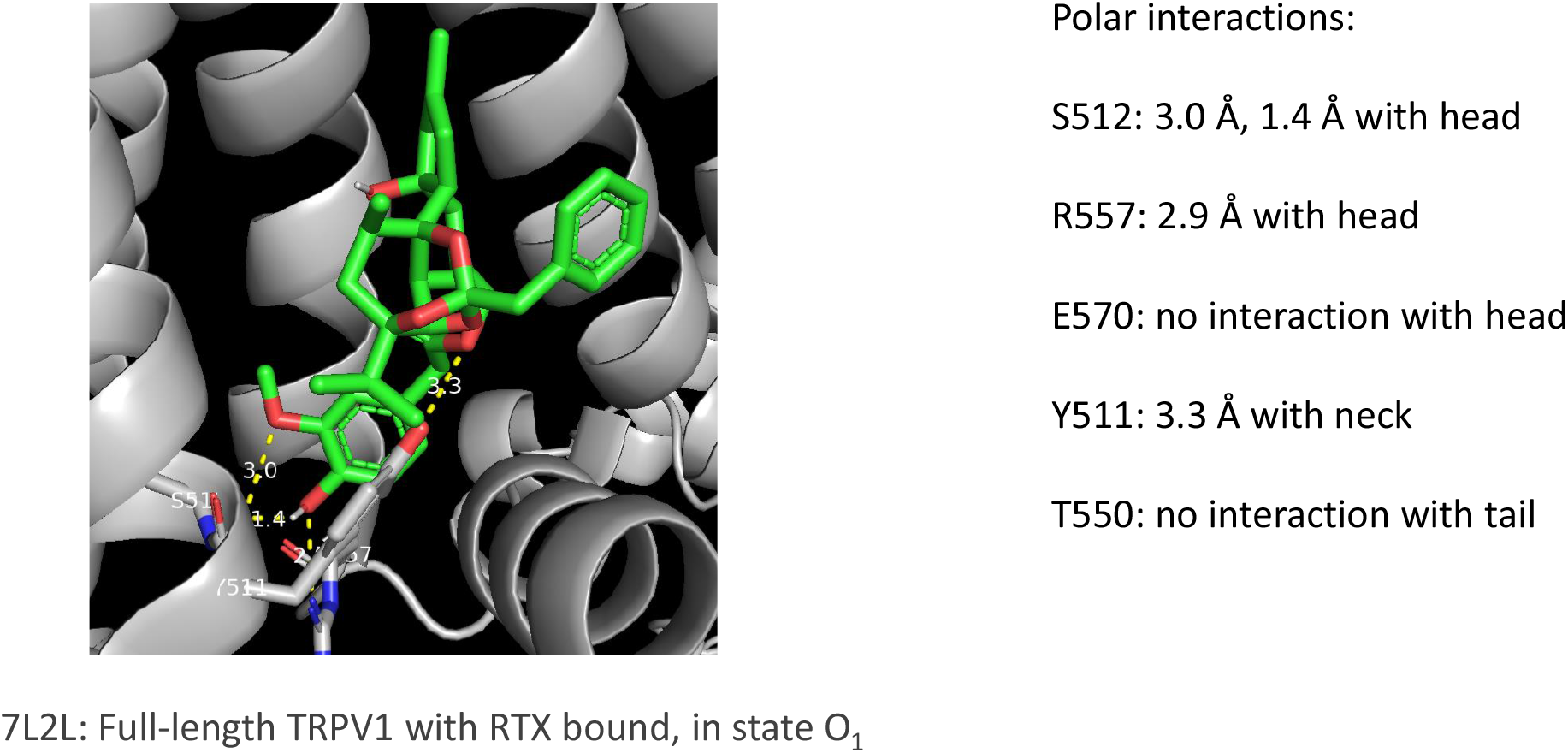

**Fig.S3.**
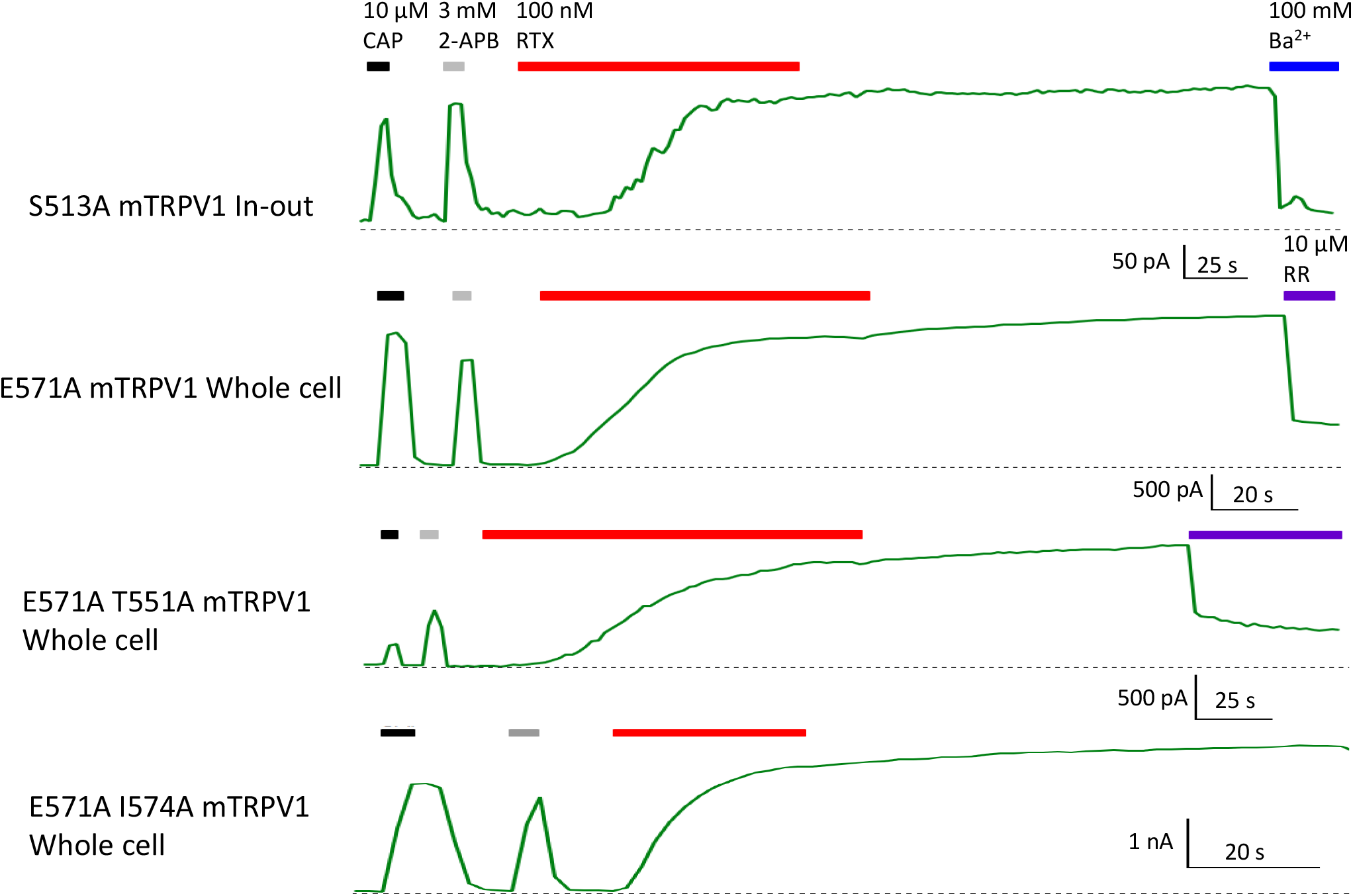

**Fig.S4.**
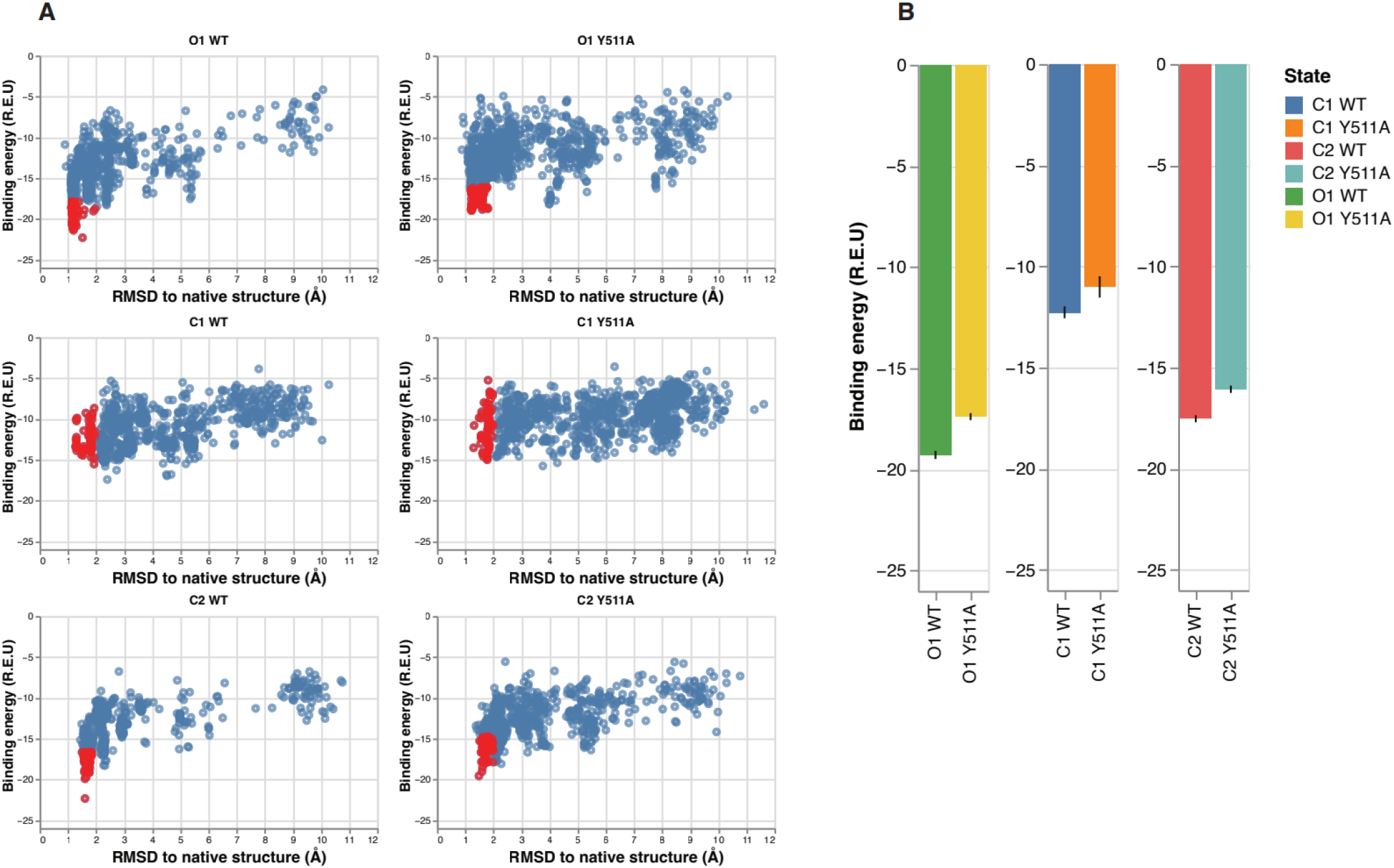

**Fig.S5.**
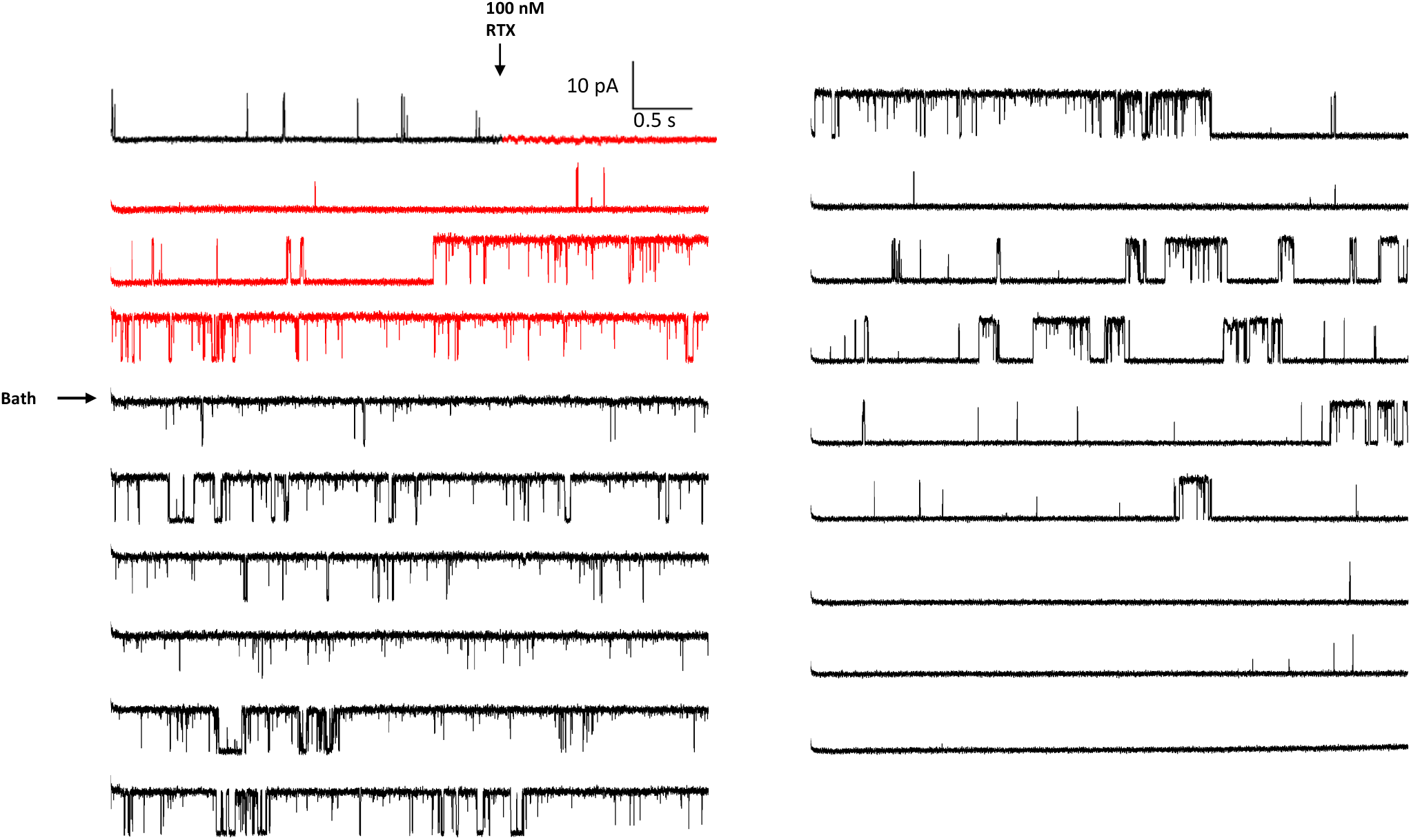
Y512A mutant

**Fig.S6.**
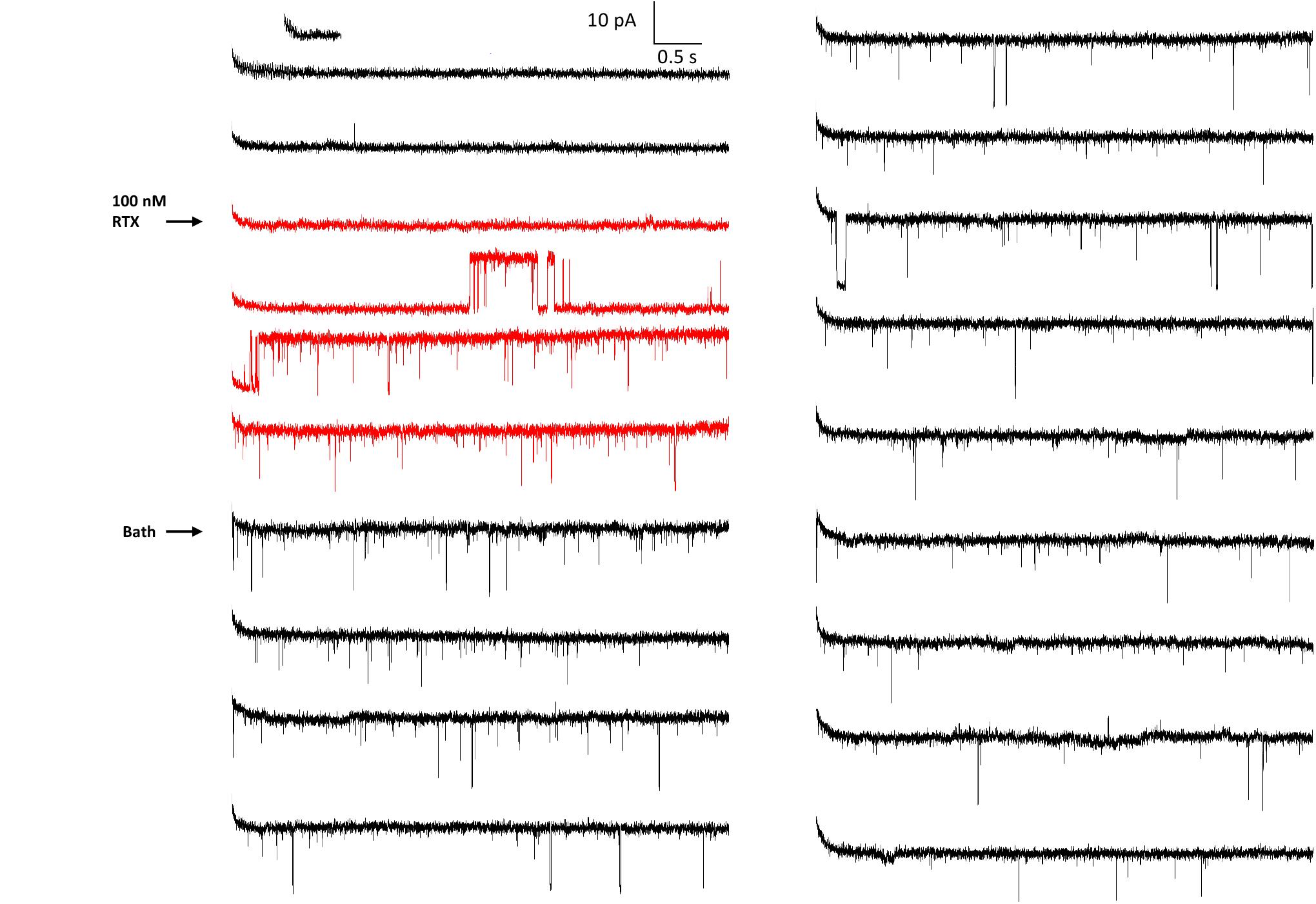
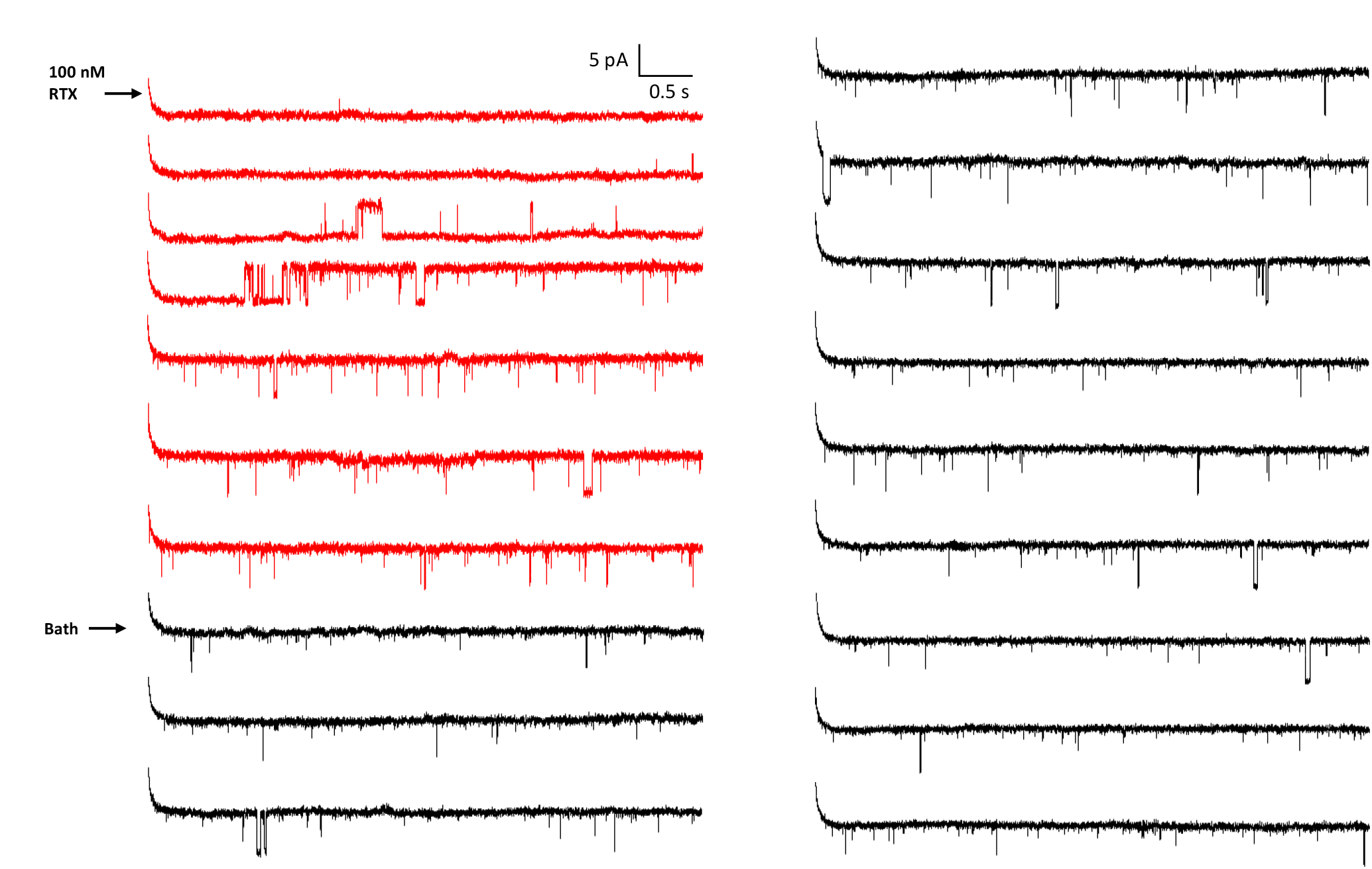
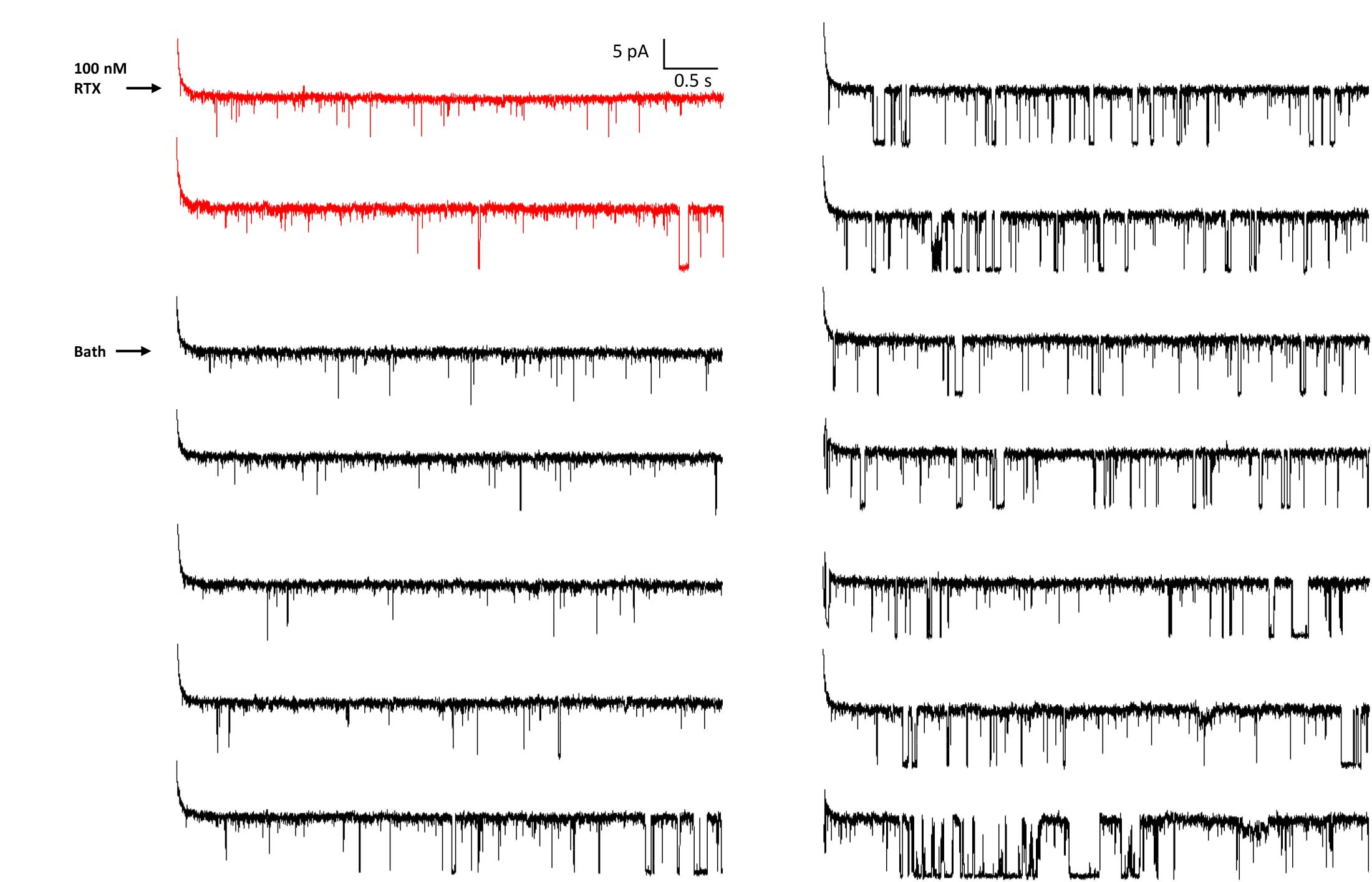
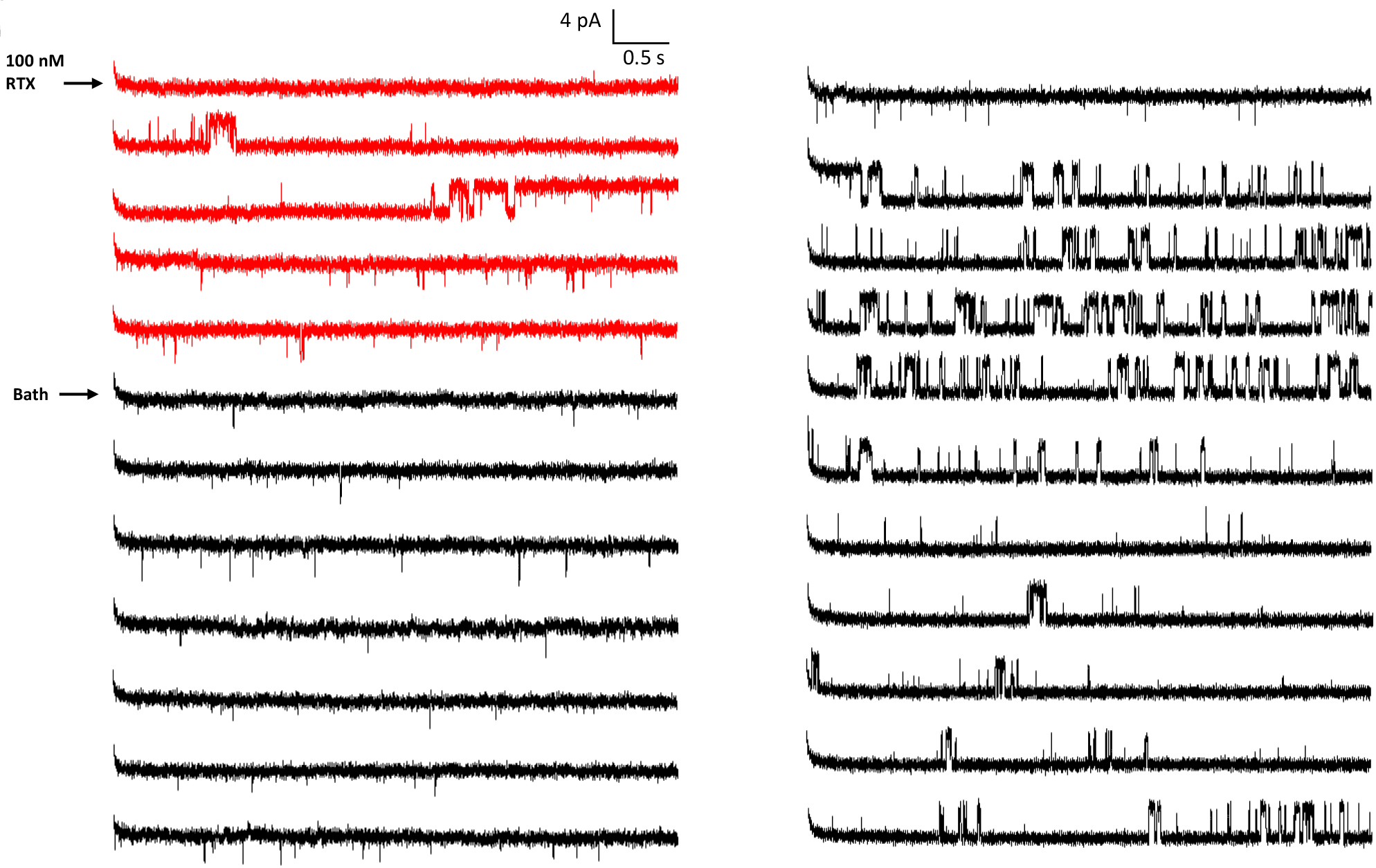
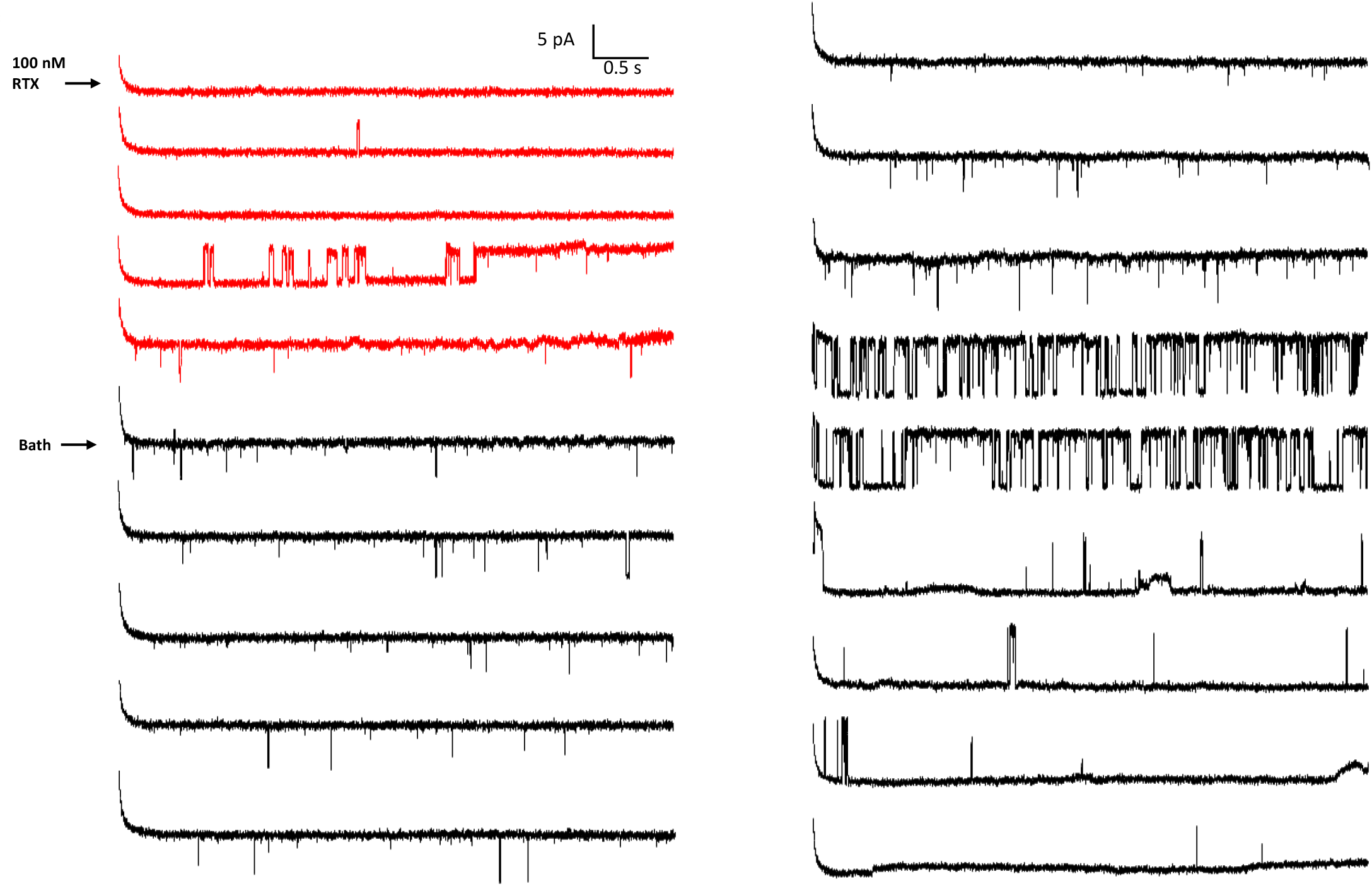

**Fig.S7.**
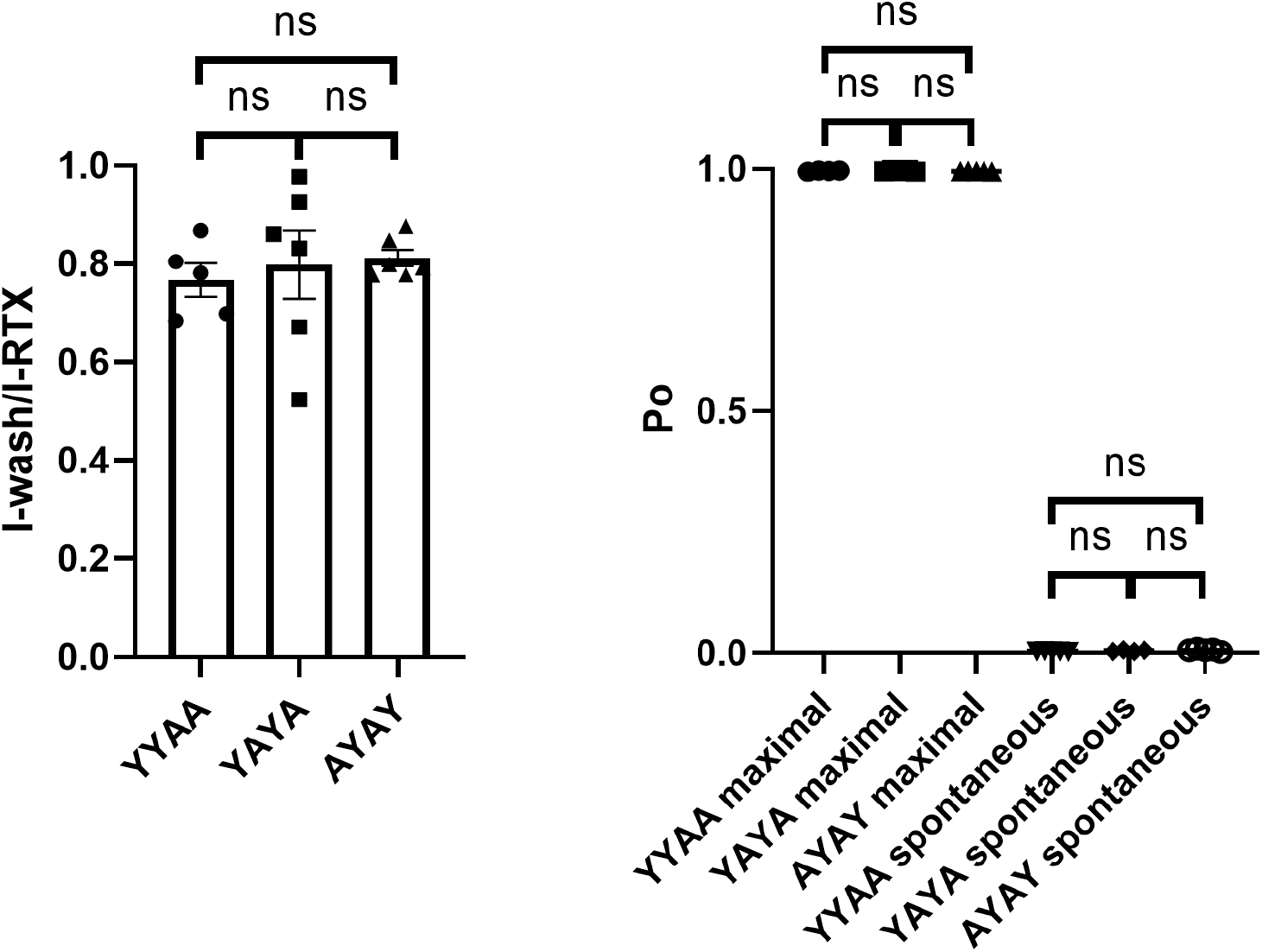
Comparison between YYAA, YAYA and AYAY concatemers

## References and Notes

1. J. Monod, F. Jacob, Teleonomic mechanisms in cellular metabolism, growth, and differentiation. Cold Spring Harb Symp Quant Biol 26, 389–401 (1961).

2. A. V. Hill, A. Paganini-Hill, The possible effects of the aggregation of the molecules of haemoglobin on its dissociation curves. The Journal of Physiology 40, 4–7 (1910).

3. J. Monod, J. Wyman, J. P. Changeux, On the Nature of Allosteric Transitions: A Plausible Model. J Mol Biol 12, 88–118 (1965).

4. D. E. Koshland Jr, G. Némethy, D. Filmer, Comparison of experimental binding data and theoretical models in proteins containing subunits. Biochemistr 5, 365–385 (1966).

5. J. N. Weiss, The Hill equation revisited: uses and misuses. The FASEB Journal 11, 835–841 (1997).

6. B. Hille, Ion channels of excitable membranes (Sinauer, Sunderland, Mass., ed. 3rd, 2001), pp. xviii, 814 p.

7. J. Zheng, M. C. Trudeau, Handbook of ion channels (Crc Press, 2015).

8. M. J. Caterina et al., The capsaicin receptor: a heat-activated ion channel in the pain pathway. Nature 389, 816–824 (1997).

9. E. Cao, M. Liao, Y. Cheng, D. Julius, TRPV1 structures in distinct conformations reveal activation mechanisms. Nature 504, 113–118 (2013).

10. R. Latorre, S. Brauchi, G. Orta, C. Zaelzer, G. Vargas, ThermoTRP channels as modular proteins with allosteric gating. Cell calcium 42, 427–438 (2007).

11. J. A. Matta, G. P. Ahern, Voltage is a partial activator of rat thermosensitive TRP channels. The Journal of physiology 585, 469–482 (2007).

12. A. Jara-Oseguera, L. D. Islas, The role of allosteric coupling on thermal activation of thermo-TRP channels. Biophys J 104, 2160–2169 (2013).

13. X. Cao, L. Ma, F. Yang, K. Wang, J. Zheng, Divalent cations potentiate TRPV1 channel by lowering the heat activation threshold. Journal of General325 Physiology 143, 75–90 (2014).

14. K. Hui, B. Liu, F. Qin, Capsaicin activation of the pain receptor, VR1:multiple open states from both partial and full binding. Biophys J 84, 2957–328 2968 (2003).

15. K. Zhang, D. Julius, Y. Cheng, Structural snapshots of TRPV1 reveal mechanism of polymodal functionality. Cell 184, 5138-5150. e5112 (2021).

16. F. Yang et al., Structural mechanism underlying capsaicin binding and activation of the TRPV1 ion channel. Nature chemical biology 11, 518–524 333 (2015).

17. S. DeLuca, K. Khar, J. Meiler, Fully Flexible Docking of Medium Sized Ligand Libraries with RosettaLigand. PLoS One 10, e0132508 (2015).

18. B. P. Brown et al., Introduction to the BioChemical Library (BCL): An Application-Based Open-Source Toolkit for Integrated Cheminformatics and Machine Learning in Computer-Aided Drug Discovery. Front Pharmacol 13, 833099 (2022).

19. A. Hazan, R. Kumar, H. Matzner, A. Priel, The pain receptor TRPV1 displays agonist-dependent activation stoichiometry. Scientific reports 5, 1–13 (2015).

20. F. Yang et al., The conformational wave in capsaicin activation of transient receptor potential vanilloid 1 ion channel. Nature communications 9, 1–9 344 (2018).

21. S. Vu, V. Singh, H. Wulff, V. Yarov-Yarovoy, J. Zheng, New capsaicin analogs as molecular rulers to define the permissive conformation of the mouse TRPV1 ligand-binding pocket. Elife 9 (2020).

22. Y. Yin et al., Structural mechanisms underlying activation of TRPV1 channels by pungent compounds in gingers. Br J Pharmacol 176, 3364–3377 (2019).

23. Y. Dong et al., A distinct structural mechanism underlies TRPV1 activation by piperine. Biochemical and Biophysical Research Communications 516, 365–372 (2019).

24. J. Schirmeyer et al., Thermodynamic profile of mutual subunit control in a heteromeric receptor. Proceedings of the National Academy of Sciences 118 (2021).

25. S. E. Gordon, W. N. Zagotta, Localization of regions affecting an allosteric transition in cyclic nucleotide-activated channels. Neuron 14, 857–864 (1995).

26. D. E. Koshland, Jr., G. Némethy, D. Filmer, Comparison of Experimental Binding Data and Theoretical Models in Proteins Containing Subunits*. Biochemistry 5, 365–385 (1966).

27. Y. Gao, E. Cao, D. Julius, Y. Cheng, TRPV1 structures in nanodiscs reveal mechanisms of ligand and lipid action. Nature 534, 347–351 (2016).

28. T. Hoshi, W. N. Zagotta, R. W. Aldrich, Biophysical and molecular mechanisms of Shaker potassium channel inactivation. Science 250, 533–538 (1990).

29. N. E. Schoppa, F. J. Sigworth, Activation of Shaker potassium channels. II. Kinetics of the V2 mutant channel. J Gen Physiol 111, 295–311 (1998).

30. J. Zheng, L. Vankataramanan, F. J. Sigworth, Hidden Markov model analysis of intermediate gating steps associated with the pore gate of shaker potassium channels. The Journal of General Physiology 118, 547–564 (2001).

31. J. Zheng, F. J. Sigworth, Selectivity Changes during Activation of Mutant Shaker Potassium Channels. Journal of General Physiology 110, 101–117 (1997).

32. D. H. Kwon, F. Zhang, J. G. Fedor, Y. Suo, S.-Y. Lee, Vanilloid-dependent TRPV1 opening trajectory from cryoEM ensemble analysis. Nature communications 13, 1–12 (2022).

33. W. Cheng, F. Yang, C. L. Takanishi, J. Zheng, Thermosensitive TRPV channel subunits coassemble into heteromeric channels with intermediate conductance and gating properties. The Journal of general physiology 129, 191–207 (2007).

34. F. Sigworth, S. Sine, Data transformations for improved display and fitting of single-channel dwell time histograms. Biophysical journal 52, 1047–1054 (1987).

