## Supplementary figures for "TRPV1 Opening is Stabilized Equally by Its Four Subunits"

**Energetics of TRPV1 Intermediate Activation States**

**Supplementary Figures**

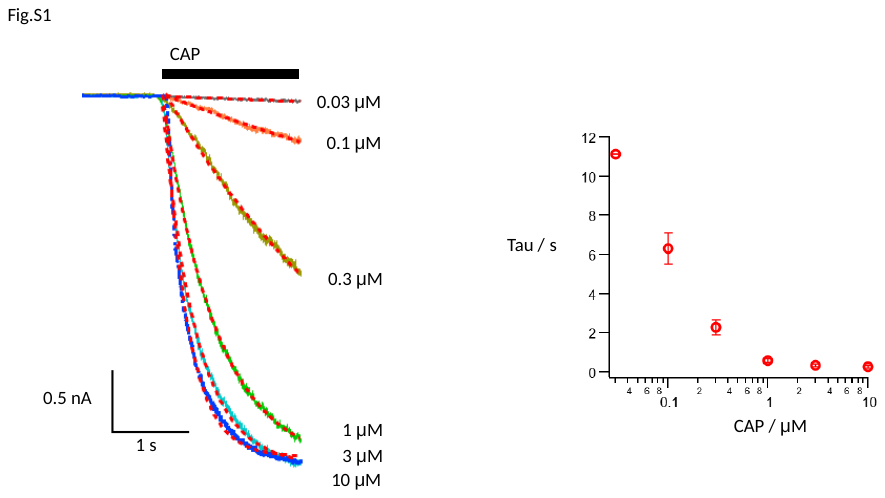

Fig. S1. Left: Representative mouse TRPV1 current traces from the same whole-cell recording activated by different concentrations of capsaicin. Superimposed are fits of an exponential function. A slight delay at the beginning of capsaicin perfusion can be seen but ignored from fitting. Right: Summarized time constant values from exponential fittings at different capsaicin concentrations, n = 6.

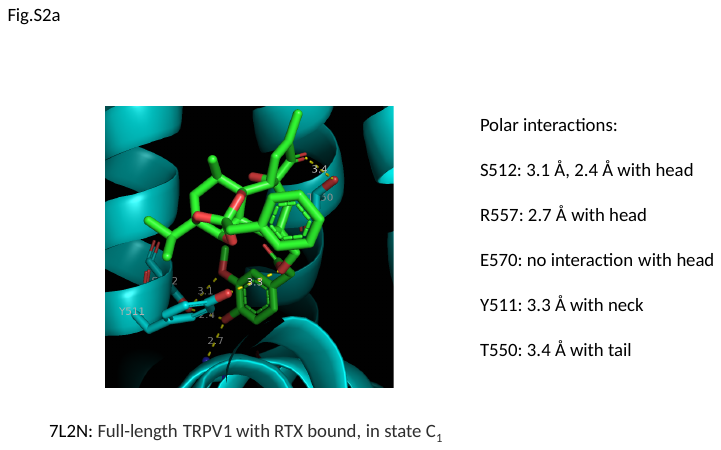

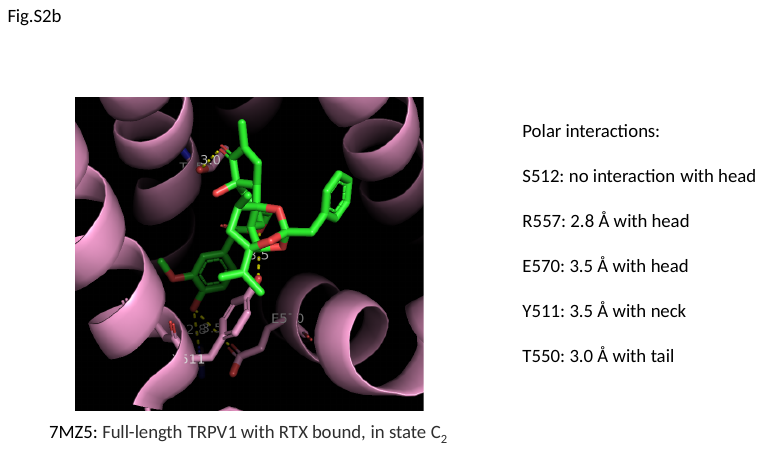

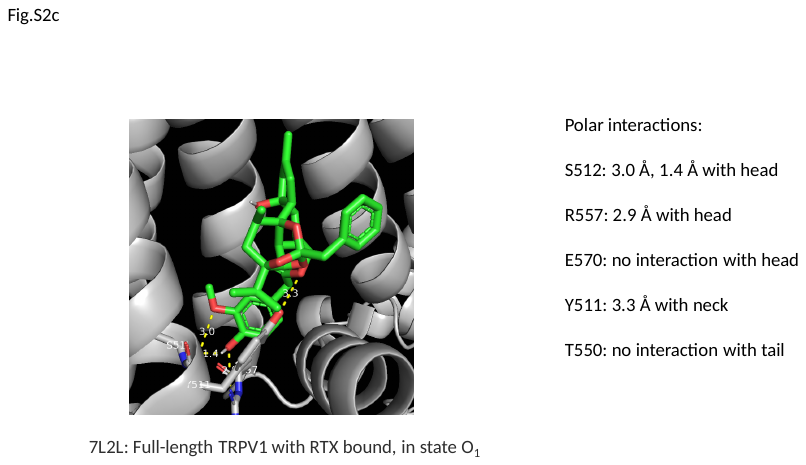

Fig. S2a-c. RTX-TRPV1 complex structures in different conformations (a, PDB: 7L2N; b, PDB: 7MZ5; c, PDB: 7L2L). The distances between RTX and potential interacting residues in TRPV1 are highlighted.

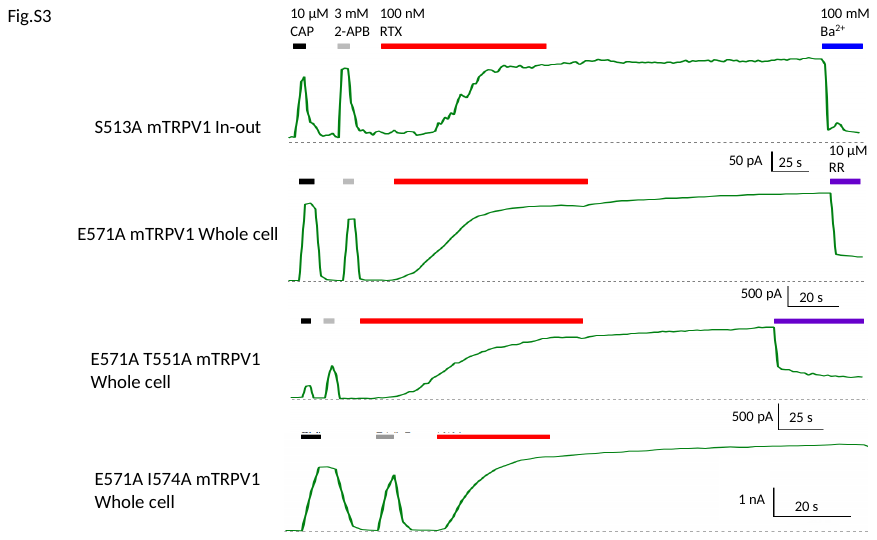

Fig. S3. Representative traces from mouse TRPV1 S513A (equivalent to S512A in rat TRPV1), E571A (equivalent to E570A in rat TRPV1), and E571A T551A double mutant (equivalent to E571A T551A in rat TRPV1) and E571A I574A double mutant (equals to E570A I573A in rat TRPV1). These mutants did not change the RTX behaviors dramatically. Representative recordings from n = 3-to-6. Ruthenium Red (RR, purple bar) or Ba^2+^ (blue bar) was used at the end of recording to block channel current and check for leak.

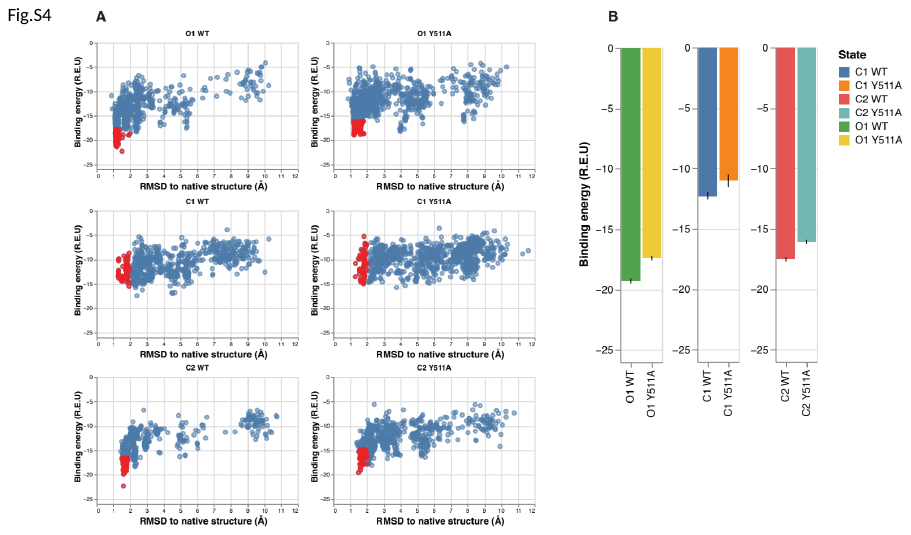

Fig. S4. RosettaLigand docking of RTX to TRPV1 wildtype and Y511A mutant structures. (A) RTX docking energy landscape is represented by the Rosetta binding energy versus the all-atom root-mean-square deviation (RMSD) to the native structure. Energy is reported in Rosetta Energy Unit (R.E.U). The data for near-native docking models (top 100 models with RMSD less than or equal to 2 Å) are shown in red. (B) Binding energy (in R.E.U.) for the top 100 near-native docking models for the wildtype (WT) and Y511A mutant in the closed (C1 and C2), and open (O1) states.

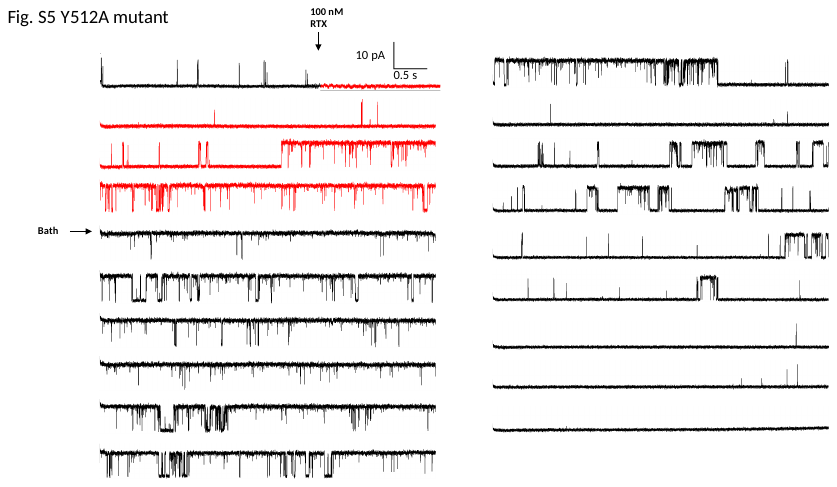

Fig. S5. Continuous single-channel recording of Y512A mTRPV1 activated by 100 nM RTX and subsequently wash-off. Events of RTX application and wash-off are indicated by arrows with current during RTX application in red. Recordings in Figure 1E originated from this recording.

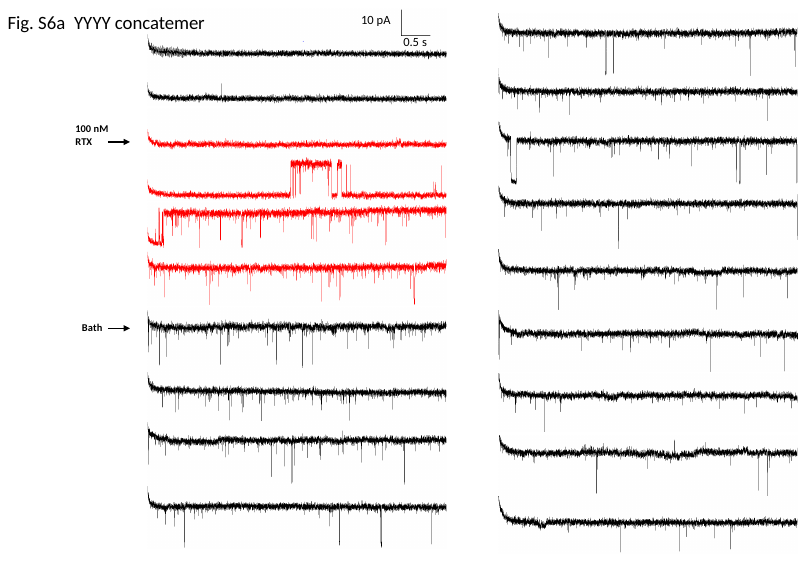

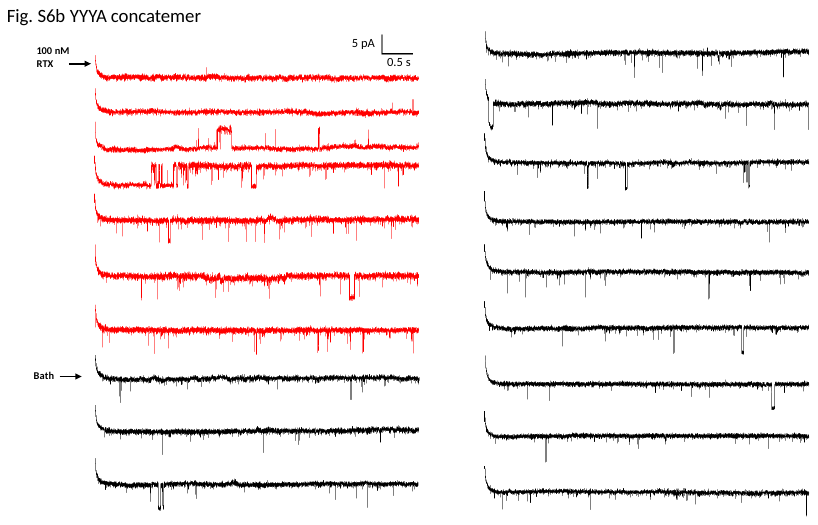

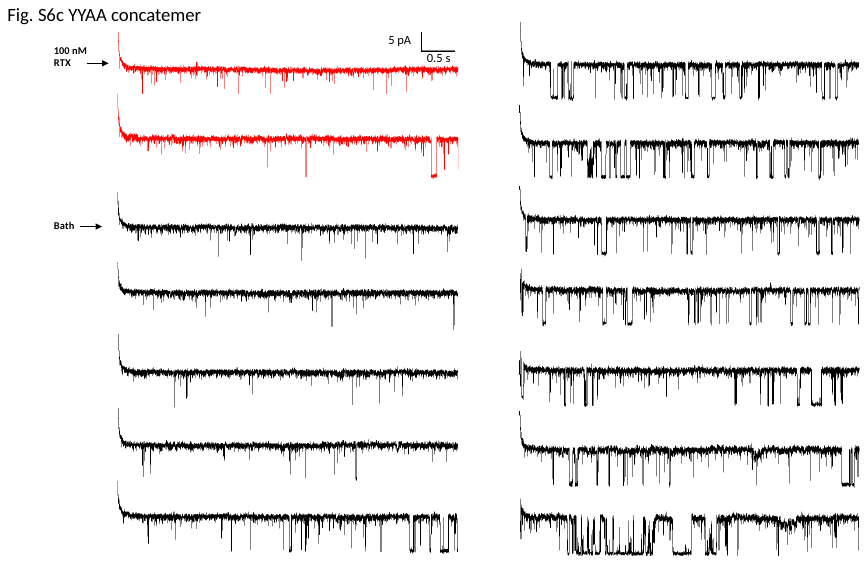

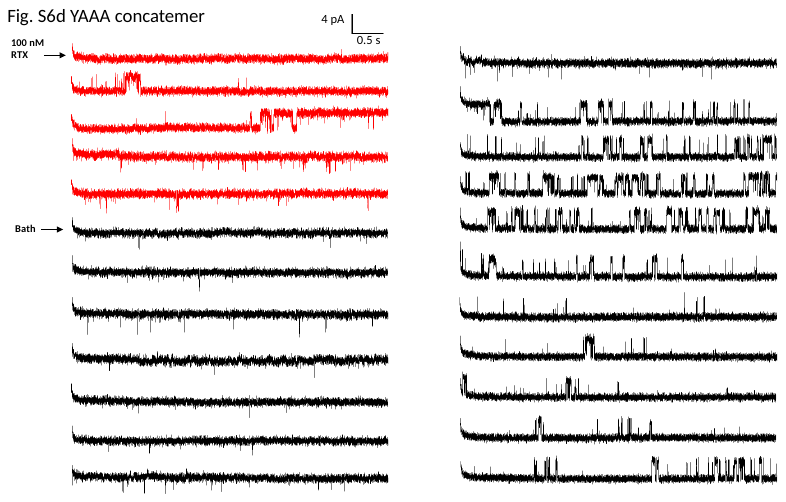

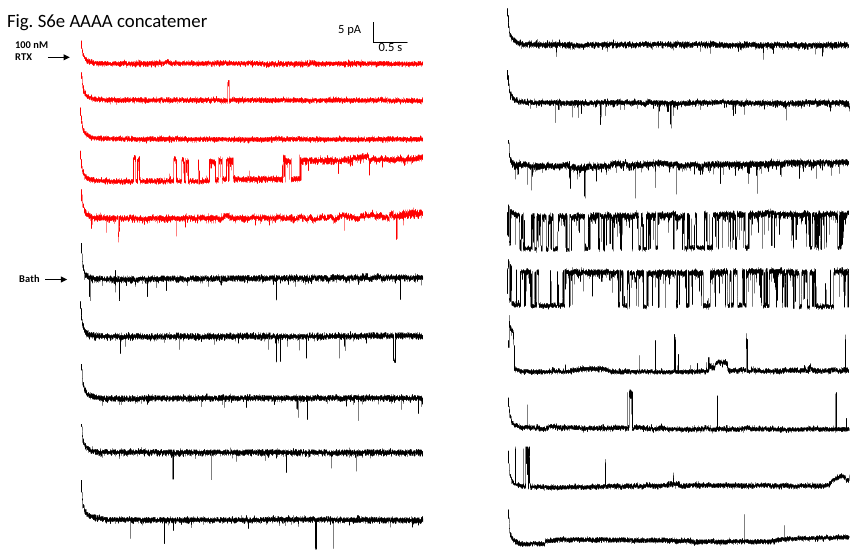

Fig. S6a-e. Representative continuous single-channel recordings of YYYY (a), YYYA (b), YYAA (c), YAAA (d), and AAAA (e) concatemer. Events of RTX application and wash-off are marked by arrows with current during RTX application in red.

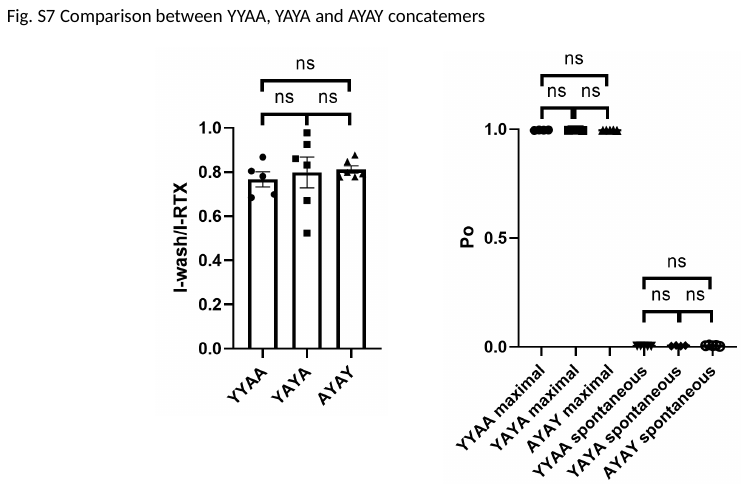

Fig. S7. Comparison between concatemers containing two wildtype and two mutant subunits with different subunit positionings. (left) Fraction of remaining current in macroscopic recordings. (right) Spontaneous and maximal open probabilities measured from single-channel recordings. n.s., not significant.
